# Spatially resolved rewiring of mitochondria-lipid droplet interactions in hepatic lipid homeostasis

**DOI:** 10.1101/2024.12.10.627730

**Authors:** Sun Woo Sophie Kang, Lauryn A Brown, Colin B Miller, Katherine M Barrows, Jihye L Golino, Constance M Cultraro, Daniel Feliciano, Mercedes B. Cornelius-Muwanuzi, Andy D Tran, Michael Kruhlak, Alexei Lobanov, Maggie Cam, Natalie Porat-Shliom

## Abstract

Hepatic lipid accumulation, or Metabolic Dysfunction-Associated Steatotic Liver Disease (MASLD), is a significant risk factor for liver cancer. Despite the rising incidence of MASLD, the underlying mechanisms of steatosis and lipotoxicity remain poorly understood. Interestingly, lipid accumulation also occurs during fasting, driven by the mobilization of adipose tissue-derived fatty acids into the liver. However, how hepatocytes adapt to increased lipid flux during nutrient deprivation and what occurs differently in MASLD is not known. To investigate the differences in lipid handling in response to nutrient deficiency and excess, we developed a novel single-cell tissue imaging (scPhenomics) technique coupled with spatial proteomics. Our investigation revealed extensive remodeling of lipid droplet (LD) and mitochondrial topology in response to dietary conditions. Notably, fasted mice exhibited extensive mitochondria-LD interactions, which were rarely observed in Western Diet (WD)-fed mice. Spatial proteomics showed an increase in PLIN5 expression, a known mediator of LD-mitochondria interaction, in response to fasting. To examine the functional role of mitochondria-LD interaction on lipid handling, we overexpressed PLIN5 variants. We found that the phosphorylation state of PLIN5 impacts its capacity to form mitochondria-LD contact sites. PLIN5 S155A promoted extensive organelle interactions, triglyceride (TG) synthesis, and LD expansion in mice fed a control diet. Conversely, PLIN5 S155E expressing cells had fewer LDs and contact sites and contained less TG. Wild-type (WT) PLIN5 overexpression in WD-fed mice reduced steatosis and improved redox state despite continued WD consumption. These findings highlight the importance of organelle interactions in lipid metabolism, revealing a critical mechanism by which hepatocytes maintain homeostasis during metabolic stress. Our study underscores the potential utility of targeting mitochondria-LD interactions for therapeutic intervention.

## Introduction

Metabolic Dysfunction-Associated Steatotic Liver Disease (MASLD) is a significant risk factor for chronic liver disease and hepatocellular carcinoma, affecting a third of the world’s population ^1, 2^. It is imperative to identify mechanisms that can reverse hepatic lipid accumulation and its associated toxicity. Interestingly, fasting also induces lipid buildup in the liver due to increased fatty acid (FA) transport from adipose tissue and a concurrent reduction in Very Low-Density Lipoprotein (VLDL) secretion ^3, 4^. Yet, how hepatocytes manage this influx of lipids during nutrient deprivation and whether these processes can be harnessed to restore function in MASLD remains unclear.

The liver plays a central role in regulating lipid metabolism, alternating between lipid oxidation and synthesis, depending on nutrient availability. Following a meal, lipids produced by the liver are sent to adipose tissue for storage. During fasting, FAs stored in adipose tissue are mobilized to the liver, where they are used to synthesize alternative fuel sources ^5^. Notably, in homeostasis, lipid metabolism is spatially segregated within the liver, with periportal hepatocytes primarily involved in lipid oxidation and pericentral hepatocytes associated with lipid synthesis ^6,7^. How these different populations of hepatocytes adapt to the influx of lipids observed during fasting and disease remains poorly understood.

Metabolic flexibility, defined as the ability to sense and respond to changes in nutrient availability, is essential for regulating adaptive organ responses ^8^. In hepatocytes, this flexibility depends on the rapid reorganization of organelles, which form interconnected networks known as metabolic sub-compartments. While the remodeling of subcellular architecture, including mitochondrial and LD morphologies, has been noted in liver physiology and disease, its functional outcomes remain unclear, partly due to the challenges of studying these processes in tissues

In this study, we examined the effects of increased hepatic lipid flux on organelle organization, focusing on mitochondria and LDs – key organelles that form contact sites to regulate lipid metabolism ^9, 10^. Using a novel approach of single-cell phenotypic profiling (scPhenomics) across different nutritional states, we discovered that mitochondria-LD contact sites are upregulated throughout the lobules of fasted mice but not in mice fed a short-term WD, to mimic the early stages of MASLD. We identified upregulation of several proteins that may facilitate this interaction, including PLIN5. We further induced the formation of mitochondria-LD contact sites through overexpression of PLIN5 and found reduced markers of lipotoxicity in WD-fed mice. These findings suggest that the dynamic assembly and disassembly of mitochondria-LD contact sites play a pivotal role in hepatic lipid management and may offer a therapeutic target for MASLD.

## Results

### Mitochondria and LD topologies mirror the spatial zonation of hepatic lipid utilization and synthesis

Hepatocytes are organized along sinusoids within the lobule, the functional unit of the liver(Fig S1A). Recent studies, including our own, have demonstrated that mitochondrial morphology, LD content, and lipid metabolism vary across the periportal-pericentral axis (PP-PC) ^11–13^. We mapped organelle topology at single-cell resolution to further investigate the relationship between organelle structure and lipid metabolism. To this aim, confocal images of liver sections from transgenic mice mtDendra2 mice ^14^ stained with BODIPY and phalloidin to mark mitochondria, LDs, and cell structure, respectively, were acquired (Fig 1A). We extracted cellular and organelle features using a custom Python-based workflow (Python v. 3.10) that employs deep-learning segmentation ^15^. The segmented hepatocytes (Fig. 1B) were categorized into bins according to their normalized distance from the central vein (Fig. 1C). The PP-PC axes were manually cropped (dashed rectangle). Similarly, we segmented organelles and extracted measurable parameters (Fig. 1D). Analysis of eighteen PP-PC axes from three mice showed an average axis length of 250 microns comprising 12-16 cells per axis (Fig. 1E). To ensure there were at least one hepatocyte per bin, we divided the PP-PC axes into 12 bins with bin 1 being the closest to the portal vessels and bin 12, closest to the central vein.

**Figure 1.**
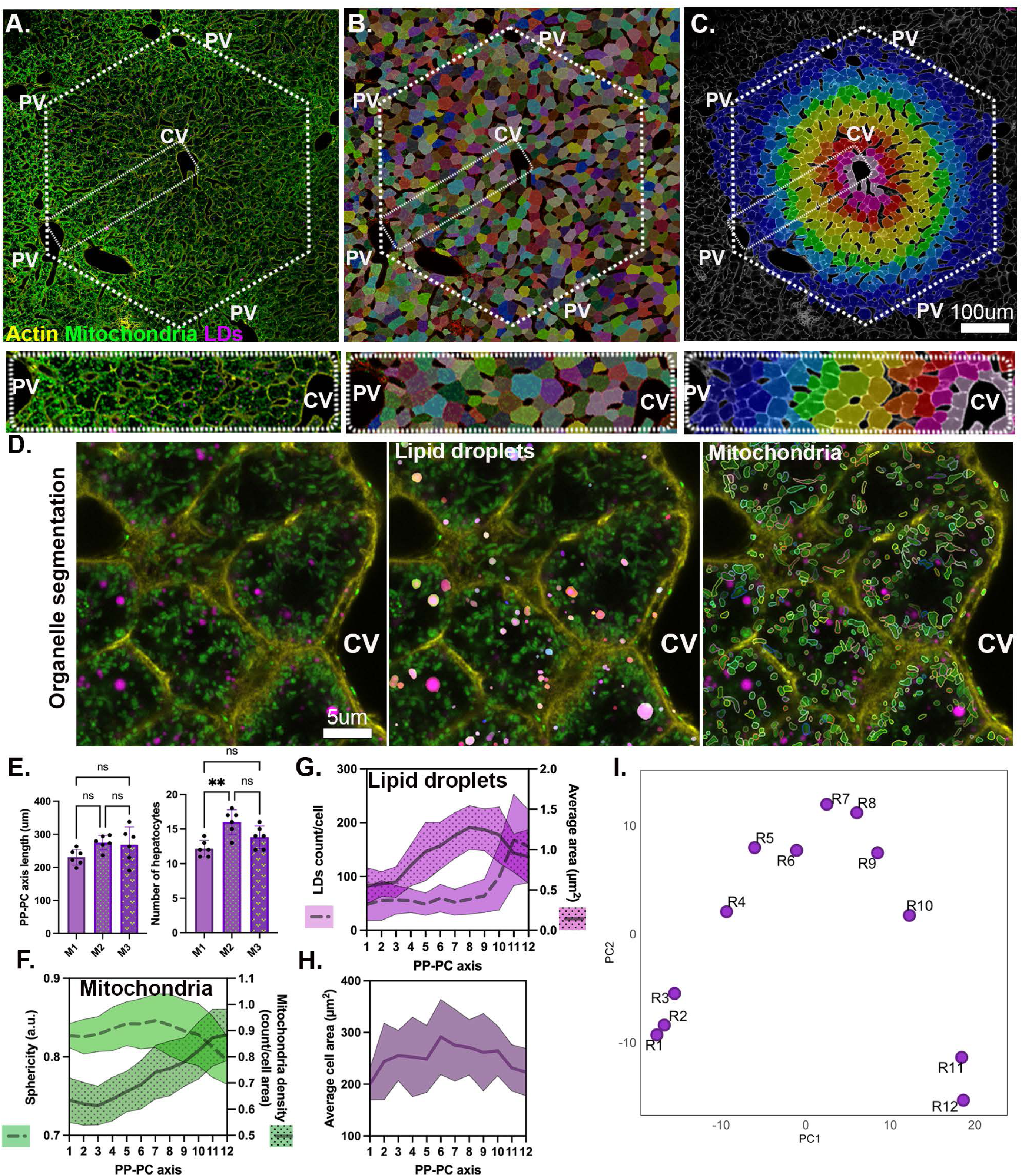
LD and mitochondrial topology define zones in the lobule. **(A)** Confocal image of a liver lobule from mtDendra2 (green) transgenic mouse. Actin was labeled with phalloidin (yellow), and lipid droplets with LipidTox (magenta). A representative PP-PC axis is marked with a dotted rectangle. **(B)** Segmentation of hepatocytes guided by actin and mitochondria labeling. **(C)** Color map representing spatial positioning of hepatocytes based on their distance from the central vein. Scale bar 100μm. **(D)** Segmentation of lipid droplets and mitochondria within cells. Scale bar 5μm**. (E)** Quantification of PP-PC axis length and number of hepatocytes per axis. Data presented as mean ± SD from 18 PP-PC axes analyzed from three mice. Statistical significance was calculated using a two-tail unpaired Student’s *t*-test). **(F-H)** Quantification of morphological features, including mitochondrial sphericity, mitochondrial density, LD count, and LD area. **(H)** Quantification of average hepatocyte cell area across PP-PC axis. **(I)** Principal Component Analysis plot of hepatocyte (R1-R12) clustering. Data presented as mean ± SD.

We analyzed the representative mitochondrial and LD features across the PP-PC axis. Mitochondria in portal and mid-lobular hepatocytes displayed higher sphericity (where a value of 1 indicates a perfect sphere) and lower density. In contrast, pericentral hepatocytes (bins 9-12) exhibited lower sphericity, indicating a more tubular morphology, along with increased mitochondrial density (Fig. 1F). LDs appeared sparse and small in the portal regions (bins 1-3). A lower LD abundance persisted through bin 9, followed by a sharp increase in bins 10-12, reflecting the higher LD content in pericentral hepatocytes (Fig. 1G). The average LD area gradually increased from bins 3-9 before slightly decreasing in pericentral regions, where LD numbers were the highest (bins 10-12; Fig. 1G).

To validate that measurements from two-dimensional (2D) sections accurately represent the three-dimensional (3D) morphology of organelles, we also assessed mitochondrial and LD features in confocal Z-stacks (Fig. S1B). We manually segmented individual hepatocytes and rendered cell and organelle surfaces using Imaris (Fig. S1B and C). The inverse relationship between mitochondrial sphericity and density that we observed in 2D was confirmed in 3D (Fig. S1D). Similarly, LD features were also zonated along the PP-PC axis in both 2D and 3D datasets (Fig. S1E). While organelle features were consistent in both 2D and 3D, the dimensionality reduction impacted cell size measurements. In the 3D dataset, PP hepatocytes were consistently smaller than PC hepatocytes within the same lobule (Fig. S1F), a trend not observed in the 2D dataset (Fig. 1H). A schematic summary illustrates mitochondrial and LD features in mice fed a control diet is shown (Fig. S1G).

Next, we examined whether topological features had specific spatial signatures. Principal Component Analysis (PCA) showed that neighboring hepatocytes cluster together (Fig. 1I). Each PCA dot represents the organelle features averaged for each bin/cell along the PP-PC axis (R1- R12). Three clusters emerged: R1-R3, R4-R10, and R11-R12. The inclusion of adjacent cells within clusters suggests that they may correspond to distinct hepatic zones specializing in lipid metabolism (Fig. 1I).

Overall, the single-cell phenotypic profiling (scPhenomics) of mitochondria and LDs efficiently used 2D datasets, allowing faster data acquisition, larger sample sizes, and automated analysis. This novel scPhenomics approach recapitulates spatial signatures associated with the functional dichotomy of lipid oxidation versus synthesis/storage, potentially defining the hepatic zones responsible for lipid handling.

### Increased lipid flux rewires organelle circuits in the zonated liver

To examine how increased hepatic lipid influx impacts organelle features, we subjected mtDendra2 mice to different dietary conditions: control diet (CNTR), fasting overnight (16 hours), or fed a WD for four weeks. Liver sections were stained with BODIPY and phalloidin to label LDs and cell structure, respectively, followed by scPhenomics workflow analysis (Fig. 2A-C).

**Figure 2.**
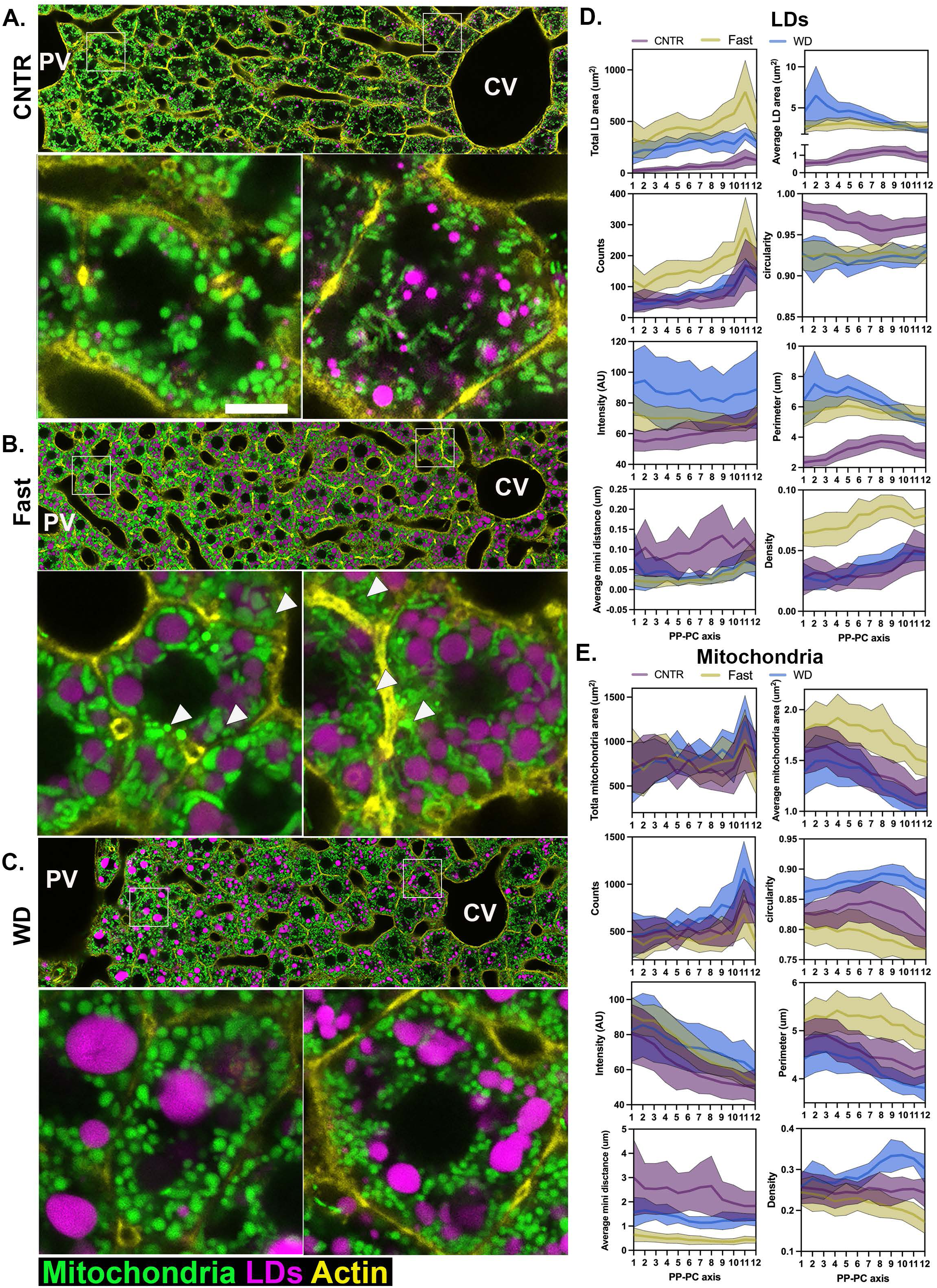
Increased lipid flux induces rewiring of LD and mitochondria circuits. **(A-C)** Representative confocal images of PP-PC axes from mtDendra2 transgenic mice fed control (CNTR) diet, overnight fasted or Western Diet (WD). Mitochondria are shown in green, actin was labeled with phalloidin (yellow), and lipid droplets were labeled with LipidTox (magenta). Per condition, representative PP and PC cells are shown in enlarged insets. Scale bar 5μm **(D-E**) LD and mitochondria features, including area (total and average per cell), count, circularity, intensity, perimeter, minimal distance, and density, are shown across space and dietary conditions. scPhenomics data is presented as mean ± SD from 18 PP-PC axes analyzed from three mice. PV: portal vein; CV: central vein; PP: periportal; PC: pericentral; LD: Lipid droplets

Fasting leads to hepatic lipid accumulation due to increased lipid influx from adipose tissue. As expected, LD content increased across the entire lobule (Fig. 2B). While LD size remained relatively uniform, the number of LDs was higher in pericentral (PC) hepatocytes. Mitochondria in periportal (PP) and mid-lobular regions appeared longer and thicker (Fig. 2B, insets). Notably, long and thick mitochondria frequently wrapped around LDs, forming extensive contact sites between the two organelles (Fig. 2B, insets). However, a subset of mitochondria within each hepatocyte retained spherical morphology and did not interact with LDs (Fig 2B arrowheads).

Dietary lipid influx also leads to hepatic lipid accumulation in WD-fed mice. Four weeks of Western diet (WD) consumption was sufficient to induce simple steatosis (Figs S2A) without causing lobular inflammation or hepatocellular ballooning to mimic the early stages of MASLD (Fig. S2B). We hypothesized that these early stages of steatosis would be particularly responsive to experimental manipulation, providing valuable mechanistic insights. Although significant LDs accumulated in WD-fed mice, the size, number, and distribution of LDs in the tissue differed markedly from those observed in fasted mice (compare Fig 2B and C). Specifically, LDs in PP and mid-lobular hepatocytes were larger than those in PC regions. The WD-fed mice had more spherical mitochondria with reduced LD interactions (Fig. 2C). Overall, increased lipid flux, whether through fasting or WD, triggered distinct remodeling of both LDs and mitochondria. Employing scPhenomics allowed for precise quantification of organelle features across different zones and dietary conditions (Fig. 2D and E). Notably, most features were influenced by both the spatial positioning of the cells and the dietary intervention. This supports the notion that LD and mitochondrial features encode spatial morphological signatures of zonation and can provide insights into the cellular metabolic state.

Next, we integrated all topological features to assess how spatial signatures changed in response to dietary perturbation. When plotted on a 2D PCA, hepatocytes from CNTR, fasted, and WD-fed mice formed tightly packed clusters, highlighting the significant impact of dietary state on organelle topology (Fig. 3A). When a third component was incorporated into the analysis, the spatial diversity within each dietary group became more pronounced. This demonstrates the distinct remodeling of organelles that depends on both the spatial positioning of the cell and the metabolic state (Fig. 3B). We used a dendrogram combined with a heatmap to further examine the information encoded by organelle features. This approach uses unsupervised clustering based on similar features. The heatmap revealed different patterns of organelle features under different nutritional conditions. CNTR-fed hepatocytes clustered into roughly three groups, consistent with the PCA data and potentially representing three zones. In contrast, fasted and WD-fed hepatocytes clustered into at least two groups of neighboring cells (Fig. 3C).

**Figure 3.**
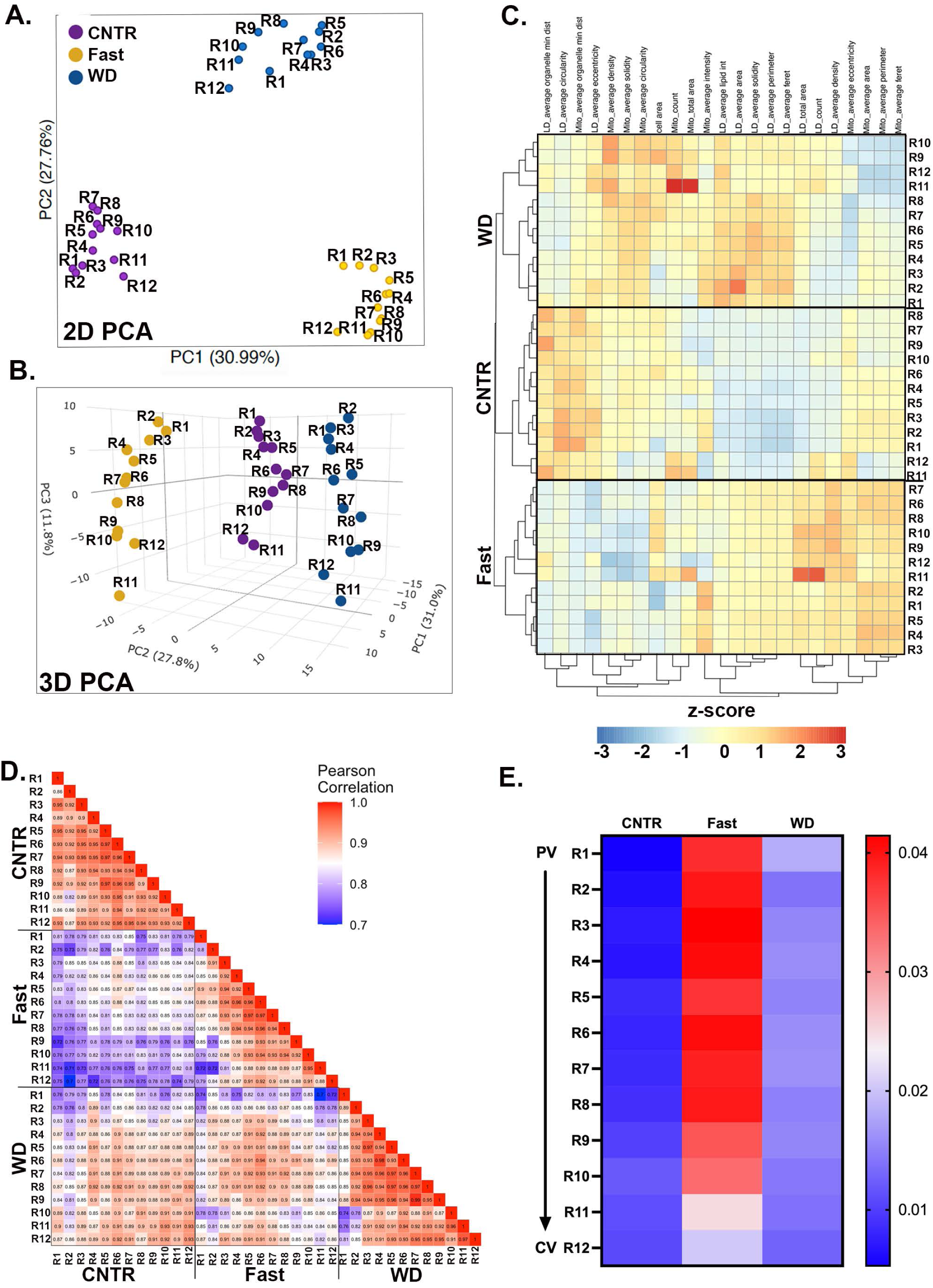
scPhenomics informs of cellular states and zonation of lipid metabolism. Principal Component Analysis (PCA) plots of single-cell organelle features (R1-R12) under different dietary conditions in **(A)** two dimensions (2D) and **(B)** three dimensions (3D). **(C)** Heatmap showing single-cell (R1-R12) organelle features under different dietary conditions. **(D)** Correlation Matrix Heatmap of organelle features under different dietary conditions. **(E)** Heat map of mitochondria and LDs overlapping pixels across the PV-CV axis (R1-R12) and diets.

To examine how individual cells respond to either fasting or WD, Pearson’s correlation coefficient was calculated, and the pairwise correlations are shown in the heatmap (Fig. 3D). Compared with CNTR cells, fasted cells (R1-R12) showed low correlation coefficient values (light-dark purple), indicating that fasting induces extensive remodeling of LD and mitochondria architecture. Conversely, most hepatocytes from WD-fed livers exhibited high correlation with the corresponding cells from the CNTR diet. However, a subset of PP hepatocytes (R1-R3) were significantly affected by WD, as demonstrated by the low coefficient values when compared with CNTR.

Of the features we measured, mitochondria-LD contacts were significantly affected by fasting and WD (Fig. 2). The distribution of mitochondria-LD contacts across the lobule and nutritional states is shown in Fig. 3E. In the CNTR diet, LD content was low and localized to PC hepatocytes (Fig 1G), consistent with the absence of mitochondria-LD overlapping pixels in PP and mid-lobular regions (R1-R8), and low pixel overlap in PC regions. Conversely, fasting dramatically increased the overlapping pixels throughout the lobule, except for R11 and R12 in PC regions. In contrast, WD-fed mice exhibited only a modest increase in overlapping pixels, with values similar to those measured in PC regions of CNTR-fed mice (Fig. 3E). The heightened LD- mitochondria interactions in response to fasting and the significant reduction of these interactions in WD-fed mice were relatively uniform across the lobule.

Next, we fasted (overnight) WD-fed mice to determine whether fasting-induced the increased mitochondria-LD interactions. We determined mitochondria-LD interactions using co- localization analysis in Imaris (Fig. S3). Fasted WD-fed mice showed notable changes in mitochondrial morphology and a significant increase in mitochondrial interactions with LDs compared to unfasted WD-fed mice. Notably, LD size increased, with larger droplets observed in fasted WD-fed mice, suggesting that mitochondria-LD contact sites may facilitate LD expansion (Fig. S3).

### Comparative proteomics of sorted hepatocytes from fed and fasted mice

To identify candidates that mediate mitochondria-LD interactions, we performed spatial proteomics. PP and PC hepatocytes were enriched from four ad-lib-fed CNTR and overnight-fasted mice using surface markers and fluorescence-activated cell sorting (FACS, as previously described ^12^). The sorted hepatocytes were subjected to tandem mass tag (TMT)-based quantitative mass spectrometry for total proteome analysis. Out of 4,999 proteins identified, 1,865 (37%) in CNTR fed mice and 1,734 (35%) in fasted mice exhibited zonation (Supplementary table). The fasting response in the different hepatocyte populations was examined using volcano plots (Fig 4A). A significant overlap was observed between PP and PC hepatocytes in both upregulated and downregulated proteins. For example, the mitophagy receptor BNIP3, is enriched in PC regions ^12, 16, 17^, was upregulated in both PP and PC hepatocytes during fasting, consistent with increased mitophagy under these conditions. Notably, over half of the PP and PC proteomes remained unchanged in response to fasting, suggesting that zonation is largely preserved during nutritional stress, with only a subset of the proteome undergoing remodeling (Fig. S4A). Pathway analysis revealed that many regulated pathways are common to PP and PC hepatocytes and were linked to mitochondrial function and lipid metabolism (Fig. S4B).

**Figure 4.**
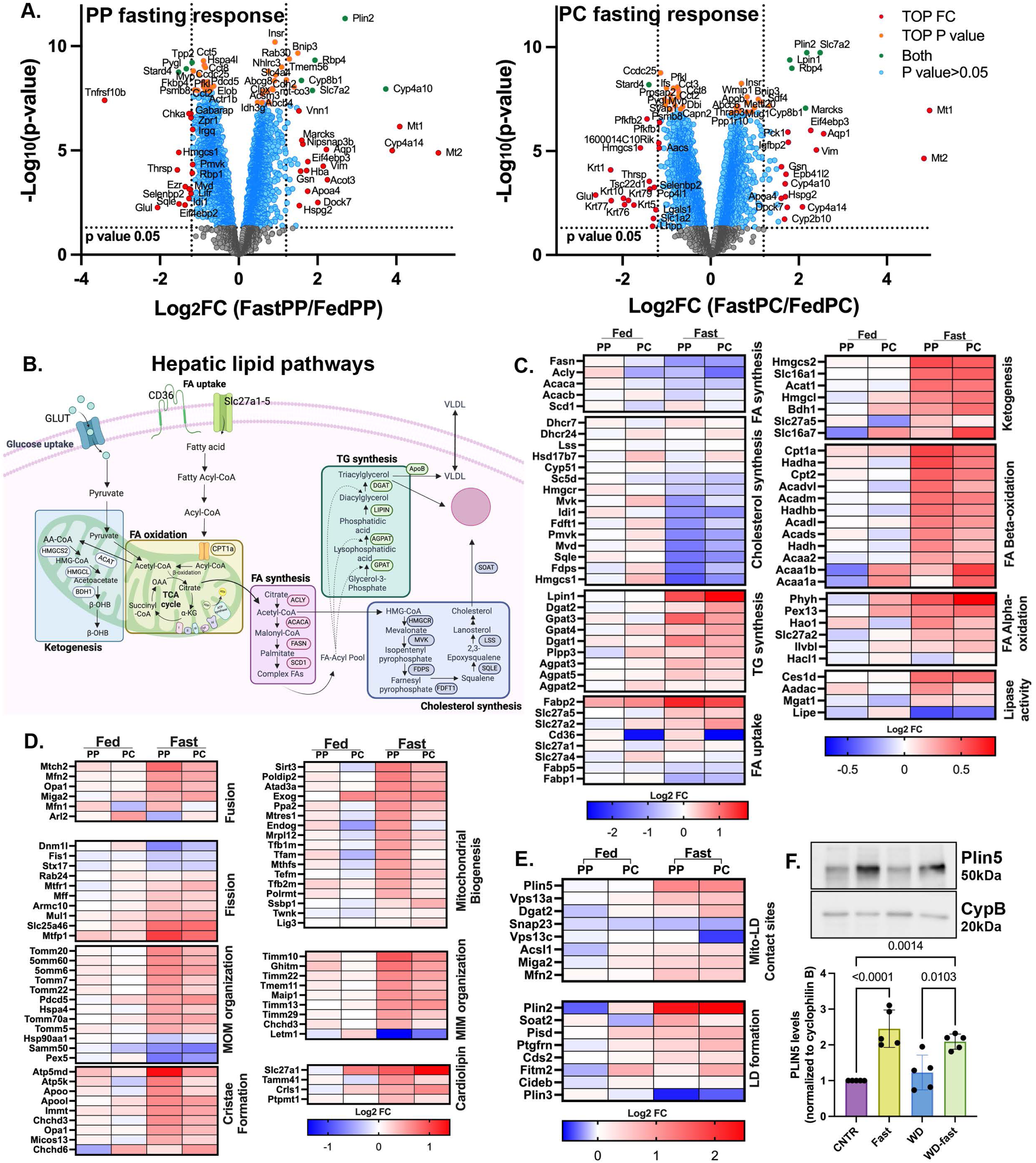
Fasting-induced remodeling of the hepatic proteome in spatially sorted hepatocytes. **(A)** Volcano plot showing the PP or PC, fast-to-fed log2 fold-change (x-axis), and the -log10 p-value (y-axis) for identified proteins. Color coding highlighting significance based on p-value and fold change. Proteomics data was analyzed with Limma R package (v3.40.6). **(B)** Schematic depicting lipid pathways in hepatocytes. **(C-E)** Heatmaps of proteins involved in lipid metabolic pathways (C), mitochondrial dynamics and structure (D), mitochondria-LD interactions, and LD formation (E) across PP and PC sorted hepatocytes from fed and fasted mice. Heatmap values were derived from the means of three independent experiments. (**F)** Western blot analysis of PLIN5 levels under different dietary conditions. Statistical significance was calculated using two-tailed unpaired Student’s *t*-test from n=5 independent experiments.

In ad-libitum CNTR diet-fed mice, many hepatic lipid pathways (summarized in Fig 4B) are zonated, as shown by the nonuniform expression of key enzymes (Fig 4C). In response to fasting, lipid pathway proteins were either upregulated or downregulated throughout the lobule (Fig. 4C compares PP and PC in the fasted state). As expected, the expression of proteins involved in FA and cholesterol synthesis was largely reduced during nutrient deprivation. An exception was observed in proteins involved in TG synthesis, an anabolic process that paradoxically increases during fasting. This likely facilitates the temporary storage of the excess free FA in LDs to prevent lipotoxicity. Additionally, proteins involved in catabolic processes, such as FA lipolysis, oxidation, and ketogenesis, were also upregulated (Fig. 4C).

In serum-deprived cultured cells, mitochondria-LD contacts are upregulated and facilitate the transfer of FA from LD into mitochondria for oxidation and ATP production ^18^. Similarly, *in vivo* fasting, mitochondria-LD contacts are formed, and proteins involved in FA uptake, oxidation, and OXPHOS are upregulated (Fig. 4C and S4C) in response to fasting. However, unlike cultured cells, mitochondrial respiration and ATP production under these conditions was significantly reduced (Fig S4D-F). During fasting, the liver shifts its metabolic focus to supplying ketones to peripheral tissues. Consistently, the levels of β-hydroxybutyrate increased more than 15-fold (Fig. S4G). These findings suggest that in the fasted liver, FAs are primarily directed to ketogenesis rather than ATP production despite the increased expression of OXPHOS proteins.

We examined the expression of proteins involved in mitochondria and LD organization and interactions. The PP-PC zonation observed in mitochondria topology and LD content corresponded to the differential expression of these proteins (Fig. 4D and E, compare PP and PC columns in fed mice). For instance, proteins associated with mitochondrial biogenesis were highly expressed in PP hepatocytes, consistent with our previous findings of greater mitochondrial mass in PP regions ^12^. Many of these proteins were upregulated in response to fasting, aligning with the extensive remodeling observed using scPhenomics (Fig. 2 and 3). Notably, PLIN5, a LD-associated protein that recruits mitochondria to form mitochondria-LD contact sites, was significantly upregulated in the fasted liver (Fig. 4E). Basal levels of PLIN5 in livers from CNTR and WD-fed mice were comparable but doubled in response to fasting (Fig. 4F).

In summary, liver zonation spatially segregates conflicting metabolic pathways across different hepatocyte populations. However, during nutrient scarcity, only essential metabolic processes are upregulated and carried out by all hepatocytes.

### Mitochondria-LD contact sites promote LD expansion in fed mice

To investigate the functional role of mitochondria-LD contact sites, we overexpressed PLIN5 in the mouse liver, using an AAV8 viral vector. Age-matched mice were fed either CNTR or WD for two weeks, followed by the overexpression of PLIN5 variants (Fig. S5). Specifically, these included the WT PLIN5, a truncated version lacking the mitochondria binding domain (PLIN5 CΔ (1-424)), and two phosphorylation mutants: a phospho-null (PLIN5 S155A) and a phospho- mimetic (PLIN5 S155E) (Fig S5). Four weeks following the viral transduction, liver tissue and serum samples were collected for microscopy and biochemical analysis (Fig. 5A).

**Figure 5.**
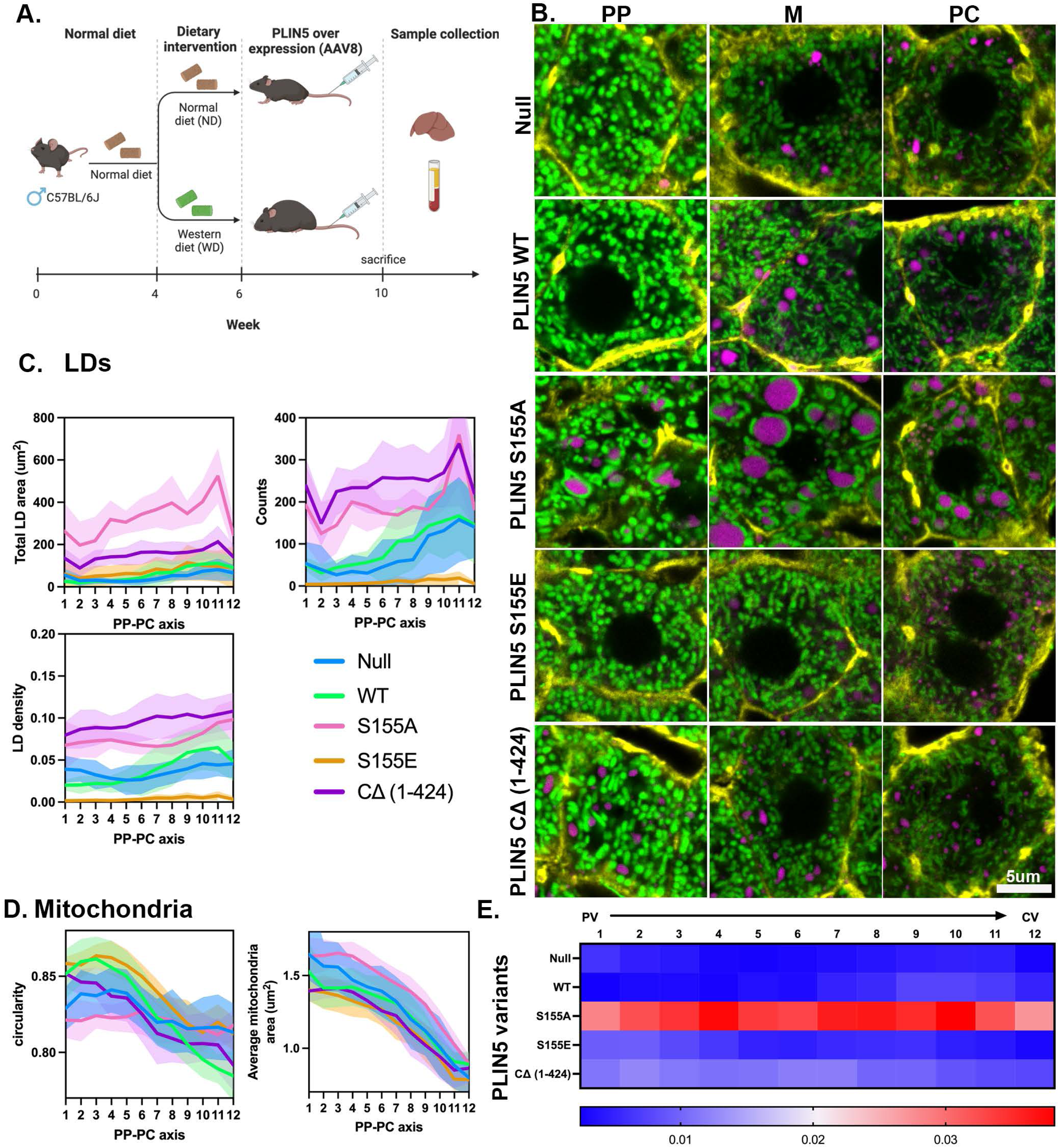
Mitochondria-LD contacts promote LD expansion. **(A)** Experimental design schematic. **(B)** Representative confocal images of periportal (PP), mid-lobular (M), and pericentral (PC) hepatocytes from mtDendra2 (green) mice fed a control diet (CNTR) and overexpressing PLIN5 variants. Actin is labeled with phalloidin (yellow), and LDs are labeled with LipidTox (magenta)Scale bar 5μm. **(C)** scPhenomics of LD features, including total area, density, and count, across the PP-PC axis from mice overexpressing PLIN5 variants. **(D)** scPhenomics of mitochondrial features, including area and circularity, across the PP-PC axis from mice overexpressing PLIN5 variants. scPhenomics data is presented as mean ± SD from 12 PP-PC axes analyzed from two mice. **(E)** Heat map of mitochondria and LDs overlapping pixels across the PV-CV axis (R1-R12) from mice overexpressing PLIN5 variants.

We assessed the impact of PLIN5 overexpression on LDs and mitochondria using scPhenomics in mice fed a CNTR diet. Representative phenotypes in PP, mid-lobular (M), and PC hepatocytes are shown in Fig. 5B (corresponding whole lobule images Fig S6A). Overexpression of WT PLIN5 caused only minor changes in LD size without altering their overall distribution. However, WT PLIN5 overexpression significantly affected mitochondrial morphology. Hepatocytes 1-6 showed an increased circularity index (spherical morphology), whereas hepatocytes 7-12 exhibited a lower circularity index (tubular; Fig. 5D comparing WT PLIN5 with the control null). Liver sections from mice expressing PLIN5 S155A revealed elongated mitochondria closely associated with large LDs throughout the lobule despite being fed the CNTR diet. In contrast, hepatocytes expressing PLIN5 S155E had fewer, smaller, and more sparsely distributed LDs (Fig 5C). Mitochondria in these cells were predominantly spherical, with a reduced mitochondria area (Fig 5D). Overexpression of the truncated PLIN5 CΔ (1-424) variant, which lacks the mitochondria binding domain, did not affect mitochondrial morphology but led to the formation of small LDs throughout the lobule, likely due to a dominant negative effect on lipolysis ^19^. Lastly, when comparing the ability of PLIN5 variants to promote mitochondria-LD interactions, PLIN5 S155A demonstrated the most pronounced effect across the entire lobule (Fig 5E). Together, these findings suggest that PLIN5 in CNTR diet-fed mice promotes TG synthesis and storage within LDs via mitochondria-LD contact sites in a phosphorylation-dependent manner.

### Overexpression of WT PLIN5 in the liver of WD-fed mice improves redox state

Next, we evaluated the impact of PLIN5 overexpression on organelle phenotypes in WD-fed mice. Representative phenotypes in PP, M, and PC hepatocytes are shown in Fig. 6A (corresponding whole lobule images Fig S7A). Overexpression of WT PLIN5 in WD-fed mice noticeably affected mitochondria length, similar to that observed in the CNTR diet-fed mice. However, under these conditions, mitochondria-LD contact sites appeared more frequent than in CNTR diet-fed mice (Fig 6A, comparing WT and null). In hepatocytes expressing PLIN5 S155A, mitochondria-LD contacts were observed throughout the lobule. In contrast, hepatocytes expressing PLIN5 S155E, displayed mitochondria that were round and that rarely wrapped around LDs (Fig 6A). In addition, the LD content in tissues expressing WT PLIN5 and PLIN5 S155E was reduced (Fig S7). As expected, the truncated variant, PLIN5 CΔ (1-424), failed to recruit mitochondria to LDs. Together, these data suggest that the overexpression of the PLIN5 S155A and S155E variants impacted mitochondrial morphology and facilitated the assembly and disassembly of mitochondria-LD contacts, respectively.

**Figure 6.**
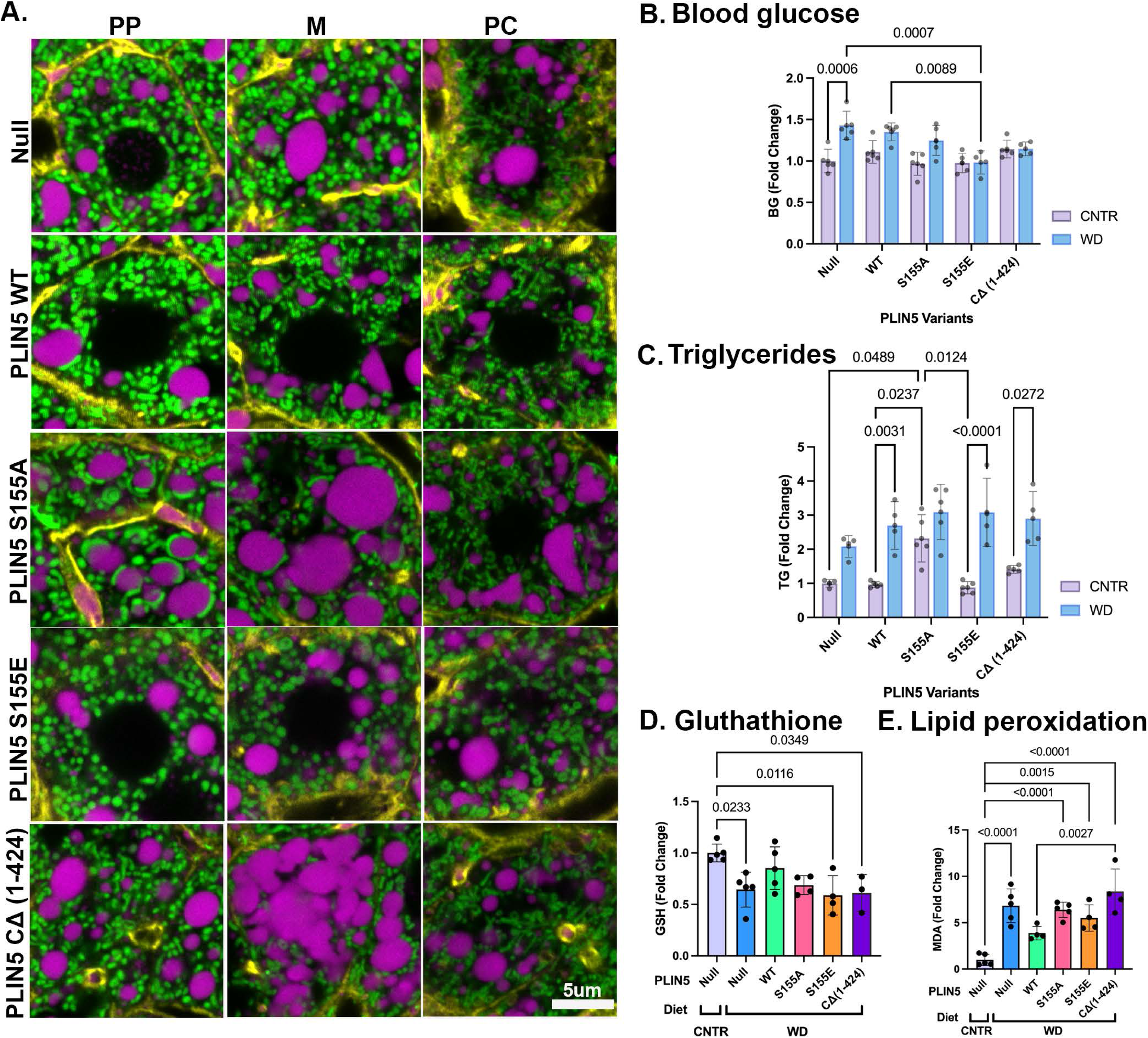
Mitochondria-LD contacts rewire lipid metabolism. **(A)** Representative confocal images of periportal (PP), mid-lobular (M), and pericentral (PC) hepatocytes from mtDendra2 (green) mice fed Western Diet (WD) and overexpressing PLIN5 variants. Actin is labeled with phalloidin (yellow), and LDs are labeled with LipidTox (magenta). Scale bar 5μm. **(B)** Blood glucose levels, **(C)** Liver triglyceride content, **(D)** Reduced glutathione levels (GSH), **(E)** Malondialdehyde (MDA) content from livers of C57BL6/J mice fed control (CNTR) or Western Diet (WD) and overexpressing PLIN5 variants. Statistical significance was calculated with two-way ANOVA. Data presented as mean ± SD from *n* = 5 independent experiments.

We then evaluated the functional consequences of mitochondrial-LD contacts to determine whether remodeling these interactions had any metabolic benefits. The increase in serum glucose levels, driven by WD, was significantly reduced in PLIN5 S155E-expressing mice (Fig 6B). In contrast, TG levels increased in PLIN5 S155A-expressing mice fed a CNTR diet, suggesting that these contact sites also promote TG synthesis. However, in WD-fed mice, TG levels did not increase beyond a certain threshold, regardless of the PLIN5 variants expressed. (Fig 6C).

Interestingly, WD-fed mice overexpressing WT PLIN5 had reduced glutathione levels comparable to those observed in mice on the CNTR diet (Fig 6D). Similarly, WT PLIN5 overexpression lowered lipid peroxidation levels in WD-fed mice (Fig 6E). The overexpression of PLIN5 variants had no effect on body weight, serum FA levels, and cholesterol (Fig S7B-D).

## Discussion

This study investigated how increased lipid flux impacts the structure, organization, and function of organelles across liver zones. We developed an imaging-based technique with a single-cell resolution to interrogate cells in their native environment within whole tissue. By mapping changes in mitochondria and LDs throughout the liver lobule, we identified distinct zonal patterns that vary depending on nutritional status. Further, we examined mitochondria-LD contact site remodeling as a potential mechanism for metabolic adaptation. These findings offer novel insights into liver zonation and organelle interactions, highlighting their role in tissue adaptation to metabolic cues. This study lends further credence to the now-recognized idea that metabolic adaptation encompasses changes to organelle structure and membrane contact sites^20^.

Distinct mitochondrial and LD characteristics were observed across liver zones in control-fed mice (Fig 1 and S1). When lipid flux was increased (via WD-fed or fasted), these organelles underwent significant adaptive remodeling, with distinct topological changes associated with WD-fed and fasted livers (Fig 2). A gold standard in liver pathology is the histological evaluation of changes to mitochondria morphology, including the emergence of megamitochondria. However, the functional implications of these changes to disease development remain poorly understood ^21, 22^. Similarly, histological evaluation of hepatic LD content is used to diagnose MASLD ^23^. Our novel system-level approach revealed spatially distinct signatures that enabled the clustering of hepatocytes into three zones (Fig 1l). These clusters shifted under WD-fed and fasting conditions, yet neighboring hepatocytes clustered together, suggesting that zonal organization remained intact during nutritional stress (Fig 3B and C). Several unique phenotypes emerged, including larger LDs in PP hepatocytes in WD-fed mice. It is likely that PP hepatocytes experience higher exposure to dietary lipids due to their proximity to the portal vein (Fig 3D). However, whether the portal buildup of larger LDs represents a protective or pathological adaptation requires further investigation. Our findings highlight the utility of scPhenomics in identifying spatially relevant phenotypes and its potential to uncover novel adaptive mechanisms driving liver disease when integrated with other single-cell methods ^13, 24^.

One prominent structural adaptation identified was the remodeling of mitochondria-LD contact sites in response to nutritional stress. In fasted livers, these contact sites spanned the lobule, while in WD-fed tissues, they were scarce (Fig 2B-C and 3E). This dynamic remodeling highlights the role of mitochondria-LD interactions in lipid metabolism, consistent with prior findings linking these contact sites to lipid oxidation or storage ^25–28^. Our findings suggest that mitochondria-LD contacts facilitate TG synthesis and LD expansion to counteract hepatic lipotoxicity in two major ways. First, fasting conditions were associated with increased LD and mitochondria-LD contact site abundance, indicating a role in lipid flux management (Fig 2B). Second, fasting in WD-fed mice restored mitochondria-LD interactions and was associated with larger LDs supporting these contacts to facilitate TG storage (Fig S3). To examine this mechanistically, we identified proteins involved in mitochondria-LD interactions using spatial proteomics (Fig 4).

Among the top upregulated proteins was the LD protein PLIN5 (Fig 4E and F). Previous studies have shown that PLIN5 recruits mitochondria to the surface of LDs ^29^. On the surface of LDs, PLIN5 inhibits lipolysis by directly binding to Adipose Triglyceride Lipase (ATGL), thus indirectly promoting LD expansion ^19^. During fasting, however, Protein Kinase A (PKA) phosphorylates PLIN5 at S155, facilitating the release of FAs via lipolysis ^9, 26, 30^. We found that *in vivo* overexpression of PLIN5 S155A, in CNTR-fed mice, promoted extensive mitochondria-LD interactions, induced TG synthesis, and increased LD size (Fig 5), supporting its proposed role in lipid storage. Conversely, overexpression of PLIN5 S155E increased lipid utilization, as evidenced by reduced TG content (Fig 6C) and significantly smaller LDs (Fig 5B). Interestingly, in starved myoblasts, the PLIN5 phosphorylation promotes the opposite function of mitochondria-LD interactions, resulting in FA transport into the mitochondria for oxidation rather than storage in LDs ^27^. These findings suggest that PLIN5-mediated mitochondria-LD contacts direct lipid flux differently depending on cell type and nutritional state. However, we did not measure the direction of FA transfer in this study due to the challenges of performing this assay *in vivo*. Further experiments are needed to fully elucidate the regulatory mechanisms across organ systems and nutritional states.

Other studies have reported similar mitochondria-LD contact site abundance increases during fasting ^3, 4^. Moreover, these contact sites have also been reported in mice fed a high-fat diet and in obese mice models ^11, 31^. However, we did not observe this increase in WD-fed mice, likely due to the shorter feeding duration (Fig S2). This aligns with the progressive nature of metabolic dysfunction in MASLD ^32, 33^, highlighting the remodeling of contact sites over time. The adaptive role of these interactions was further supported by findings that overexpression of WT PLIN5 improved markers of lipid peroxidation and redox state (Fig 6D and E). Other studies have demonstrated that increasing PLIN5 levels may benefit glycemic, lipid control, and overall cellular health ^29, 34, 35^. We propose that increased lipid influx during fasting remodels mitochondria-LD contact sites, promoting TG synthesis and preventing adverse lipotoxicity. In contrast, lipid influx resulting from nutrient excess, as seen in the early stages of MASLD, leads to harmful effects due to inappropriate organelle remodeling, elevated free fatty acids that are improperly stored, and oxidative stress. Notably, the overexpression of WT PLIN5 in Western diet (WD)-fed mice alleviates some of this damage by enabling cells to better regulate organelle interactions. These findings suggest that dynamic contact sites are critical in efficient lipid handling and maintaining cellular health, highlighting potential therapeutic targets for metabolic disorders (Fig. 7).

**Figure 7.**
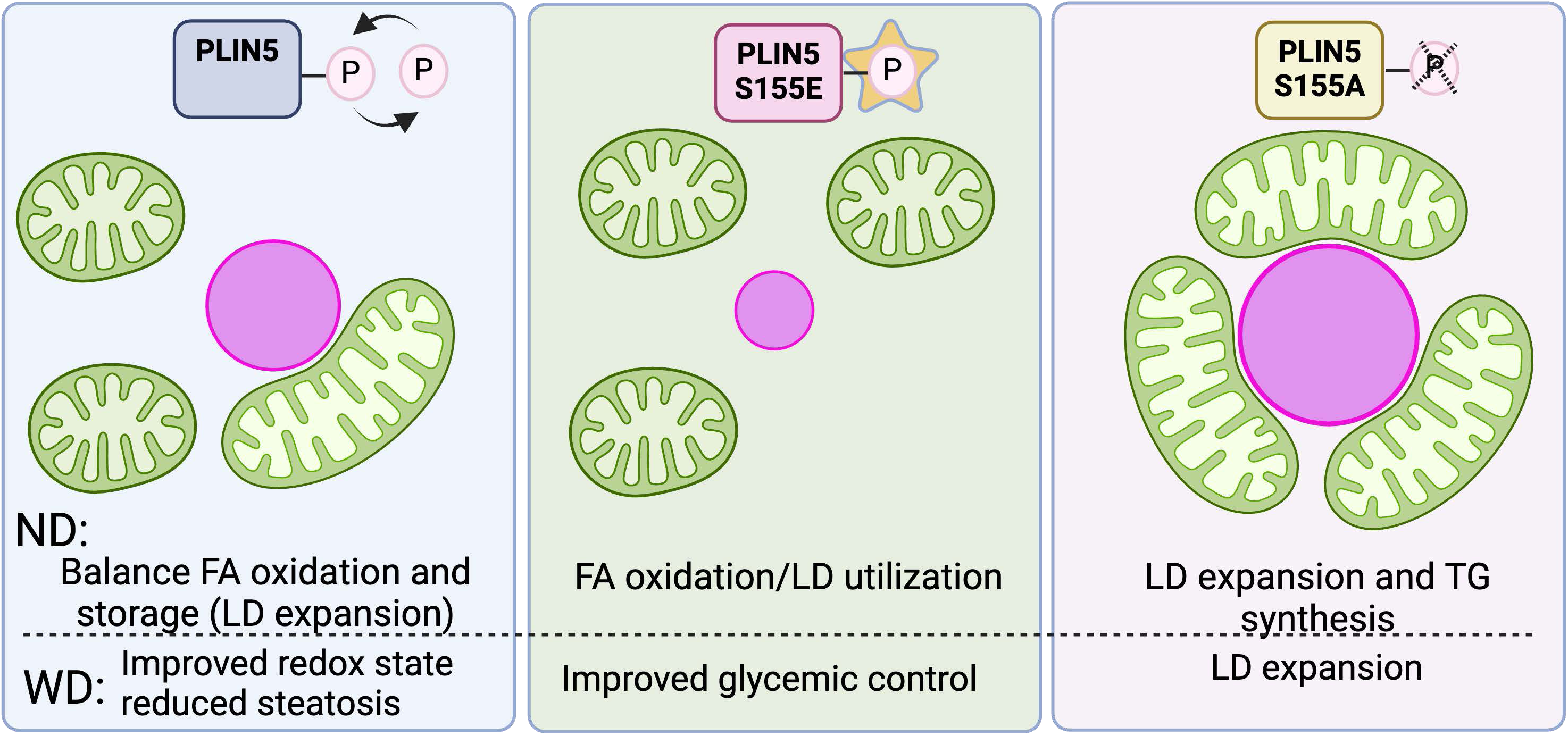
Regulation of lipid metabolism via PLIN5-induced mitochondria-LD contacts. Proposed model of mitochondria-LD contacts regulation by PLIN5. PLIN5 induces mitochondria-LD contacts in a phosphorylation-dependent manner. PLIN5 S155A induces contact sites promoting TG synthesis and storage in LD. On the other hand, in PLIN5 S155E, mitochondria and LD are rarely associated, and LDs are smaller, indicating FA oxidation. In WD-fed mice, WT PLIN5 improves cellular health by reducing oxidative stress. The model was created with Biorender.com.

The Endoplasmic Reticulum (ER), in conjunction with mitochondria and LDs, plays a central role in lipid synthesis and metabolism ^11, 36, 37^. We identified proteins involved in mitochondria-LD contacts, such as Mitofusin 2 (Mfn2) and Mitoguardin 2 (MIGA2), which localize to the ER and coordinate lipid synthesis (Fig 4A). These interactions link mitochondrial lipid supply to TG production in the ER and its subsequent storage in LDs ^38^. Given the central role of the ER in glycemic control and lipid metabolism, the observed metabolic benefits of PLIN5 overexpression in WD-fed mice may, in part, result from ER-mediated processes (Fig 6D and E). scPhenomics could further elucidate these interactions by expanding the analysis to additional organelles, as demonstrated in cell culture studies ^25, 39^.

Our findings highlight the significance of mitochondrial morphology and dynamic organelle interactions in regulating metabolic flexibility. In the fasted state, most mitochondria are tubular and closely associated with LDs, though a subset of spherical cytosolic mitochondria is also present (Fig. 3B). Previous studies have identified distinct mitochondrial subpopulations within tissues, with mitochondria associated with LDs being metabolically distinct from their cytosolic counterparts ^26, 28, 40^. In homeostasis, mitochondria in PP hepatocytes are oxidizing FA (cytosolic with round morphology), while those in PC hepatocytes are synthesizing TGs and storing them through contact with LDs ^12^. However, during fasting, the prominence of distinct mitochondrial subtypes may increase. This could explain how, under nutritional deficits, mitochondria simultaneously support lipid oxidation for ketogenesis while promoting TG synthesis and storage to mitigate lipotoxicity. In WD conditions, PLIN5 overexpression enables hepatocytes to regulate the proportion of mitochondria bound to LDs versus those free in the cytosol, facilitating more efficient management of lipid overload (Fig. 7). PLIN5-mediated reorganization of mitochondria-LD interactions highlights a key metabolic adaptation and suggests potential therapeutic strategies for metabolic disorders such as MASLD. Examining mitochondria-LD interactions in the zonated liver may also play a role in hepatocellular carcinoma^16, 41^.

Overall, our study highlights the complexity of liver zonation and the adaptive role of organelle interactions under varying metabolic conditions. Future studies expanding scPhenomics to additional organelles could deepen our understanding of inter-organellar organization in liver physiology and disease. This approach provides a powerful platform for dissecting the spatial dynamics of lipid metabolism and their implications in disorders such as MASLD.

## Contact information

Natalie Porat-Shliom, Center for Cancer Research, National Cancer Institute. Building 10, Room 12C207 MSC 1919, Bethesda, MD 20892. Phone: 240-760-6847.

## Acknowledgments

We thank Dr. Sarah Cohen, UNC-Chapel Hill, for the PLIN5 constructs. We thank Drs. Sudipto Das and Thorkell Andresson from the CCR proteomics Facility at the Frederick National Laboratory for Cancer Research. We thank Pranali Pathare Mangat of 3P Scientific Communications for scientific editing.

---

**Supplement Figure 1.**
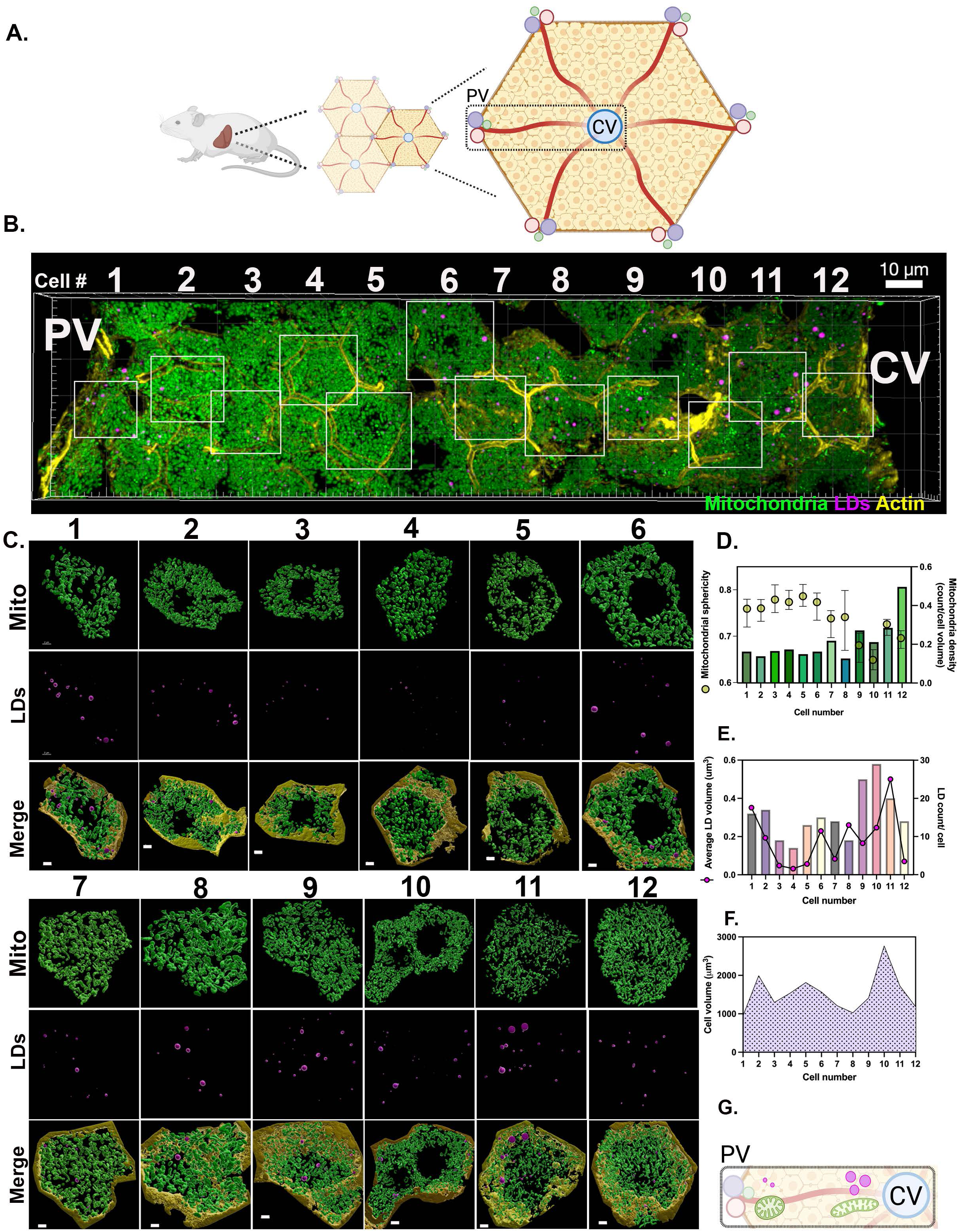
Single-cell phenotyping of LD and mitochondrial topology in 3D. **(A)** Anatomical organization of the mouse liver lobule. The model was created with Biorender.com. **(B)** Representative projection of z-stack from mtDendra2 mouse visualizing mitochondria (green), lipid droplets (magenta), and actin (yellow). Scale bar 10 μm. **(C)** 3D surfaces of mitochondria (green), LDs (magenta), and actin (yellow) at a single-cell resolution. Cells are numbered from 1 to 12 on the PV-CV axis. Scale bar 2 μm **(D)** Quantification of mitochondrial sphericity and density. **(E)** Quantification of lipid droplet volume and count. **(F)** Quantification of cell volume. PV: portal vein; CV: central vein; LD: lipid droplet.

**Supplement Figure 2.**
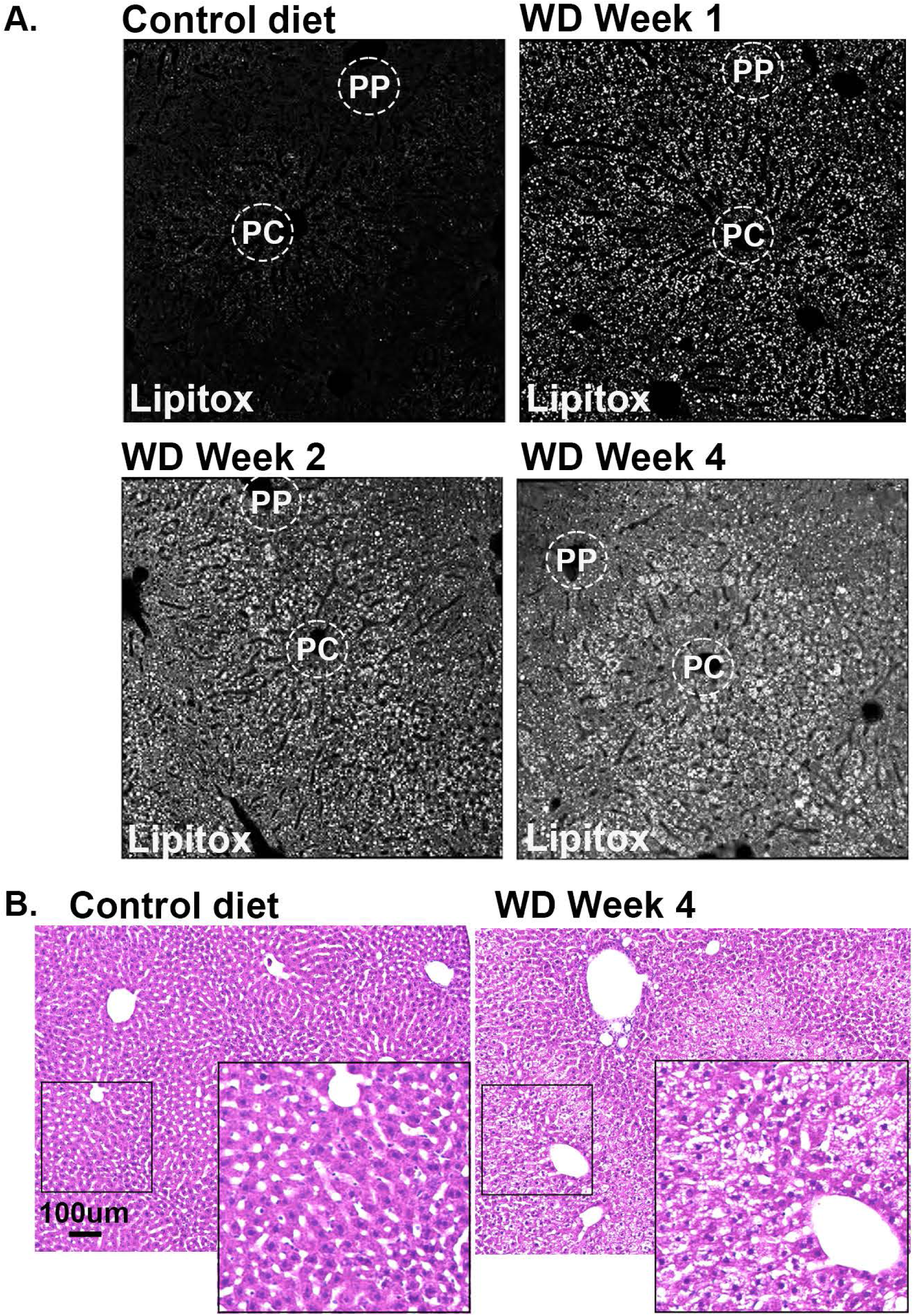
Lipid droplet accumulation in livers from mice fed WD for 4 weeks. **(A)** Confocal images of liver sections from C57BL6/J mice fed a control diet or a Western Diet as indicated. Lipids were labeled with LipidTox (white). **(B)** H&E staining images of liver sections from mice fed with control or western diet for 4 weeks. Scale bar 100 μm PP: periportal; PC: pericentral; WD: western diet.

**Supplement Figure 3.**
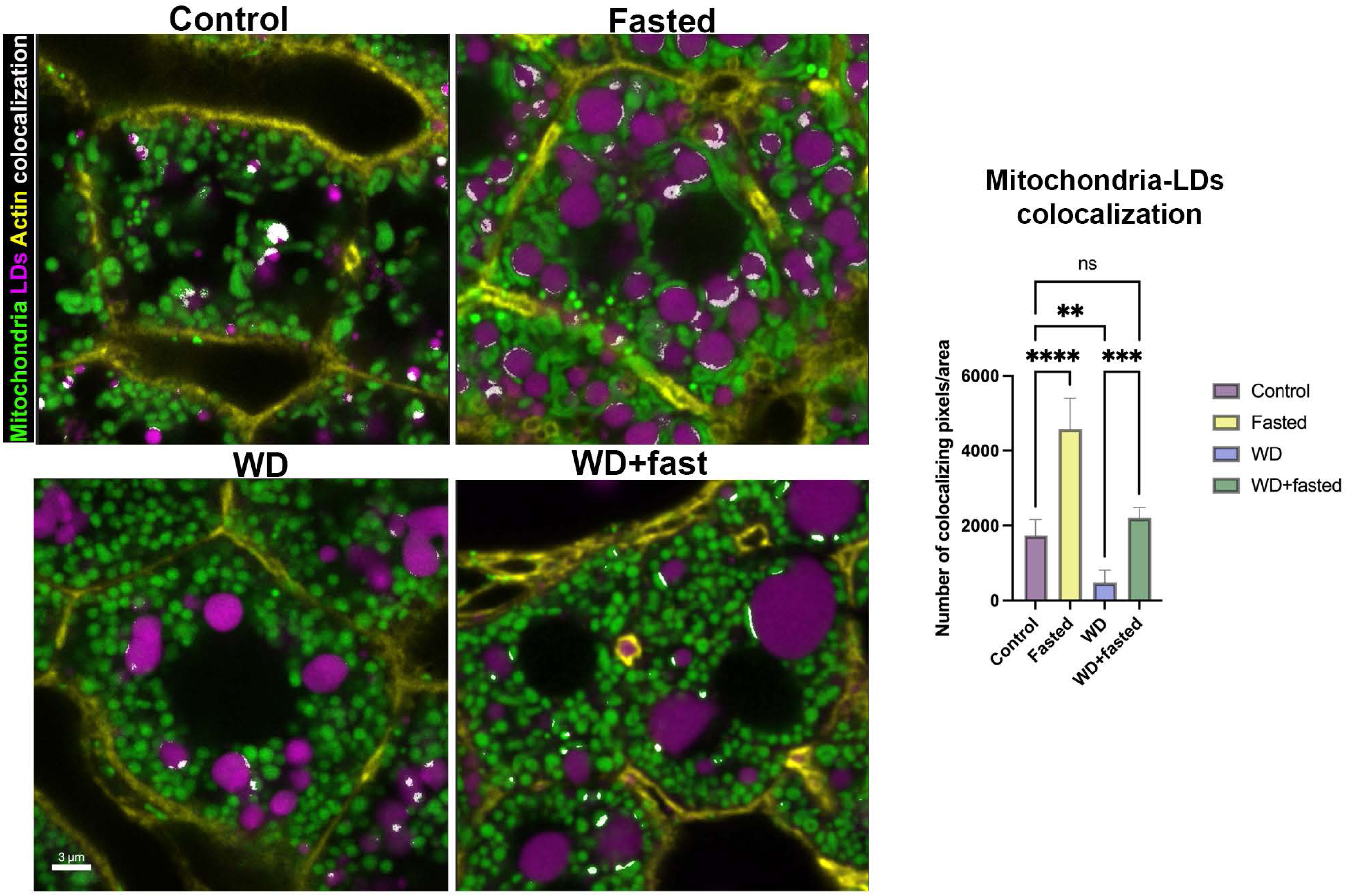
Fasting induces Mitochondria-LD interactions in WD-fed mice. **(A)** Confocal images of liver sections from mtDendra2 (green) transgenic mice fed control, WD, overnight fasted or WD-overnight fasted. LDs were labeled with Lipidtox (magenta) and actin with phalloidin (yellow). Colocalization between mitochondria and LD is shown in white. Scale bar 3μm. **(B)** Quantification of mitochondria-LD colocalization under different dietary conditions. Statistical significance was calculated using two-tailed unpaired Student’s *t*-test. Data presented as mean ± SD from *n* = 5 independent experiments. \**p* < 0.05, \*\**p* < 0.01, \*\*\**p* < 0.001, \*\*\*\**p* < 0.0001. WD: western diet; LD: lipid droplets.

**Supplement Figure 4.**
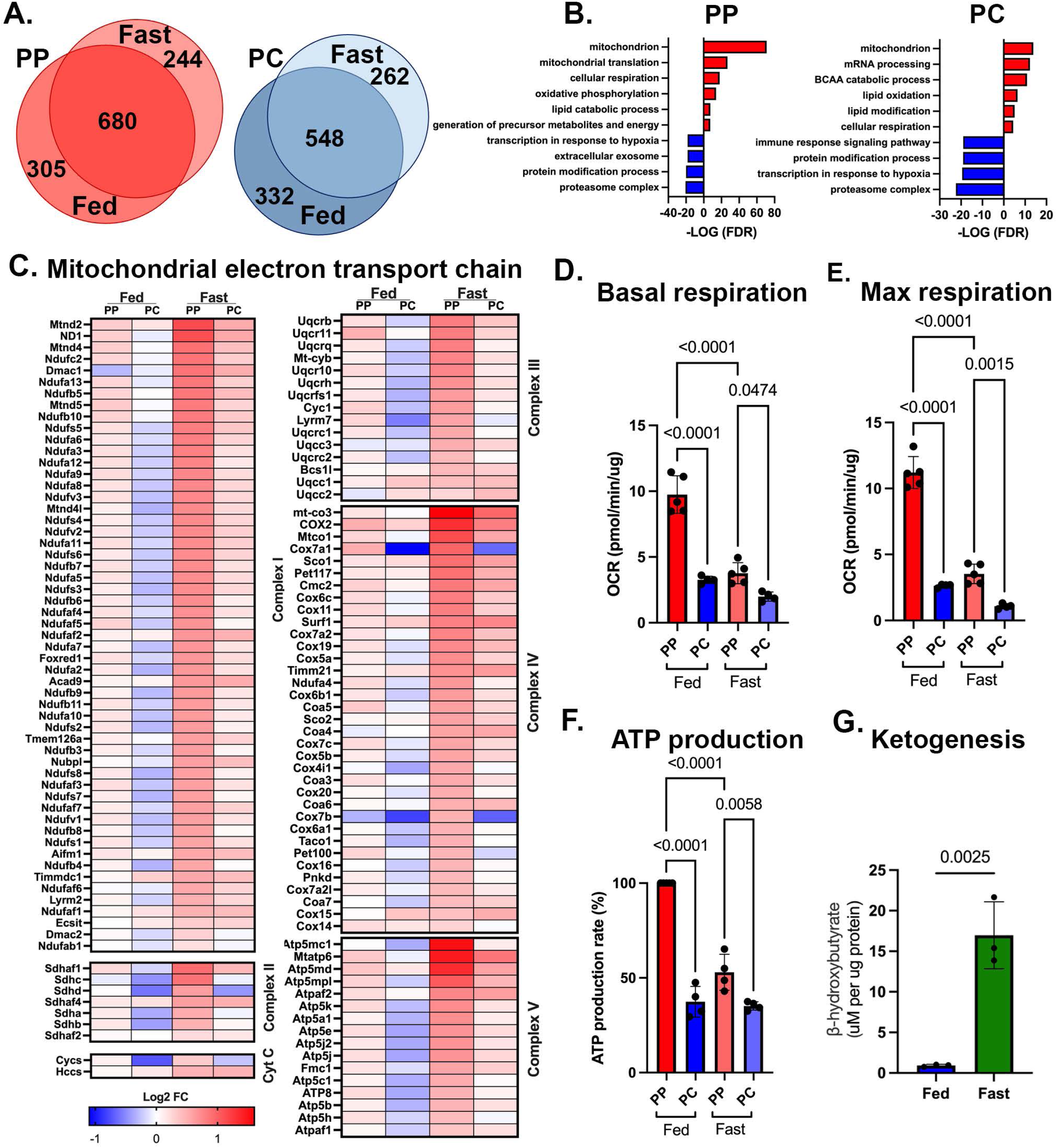
Fatty acids drive ketogenesis, not OXPHOS, during fasting. **(A)** Venn diagram depicting PP or PC signature proteins in fed and fasted mice. **(B)** GO enrichment analysis for the proteomics data in the spatially sorted cell. **(C)** Heatmap showing levels of proteins in the mitochondrial respiratory complex in spatially sorted hepatocytes from fed and fasted mice. Values were derived from the means of three independent experiments. Basal and maximum respiration **(D-E)** and ATP production **(F)** in spatially sorted hepatocytes. Oxygen consumption rates (OCR) were measured with XF Mito Stress Test Kit, and Seahorse XF96 Analyzer. Samples were normalized to cell number. Statistical significance was calculated with two-way ANOVA. Data presented as mean ± SD from *n* = 4 independent experiments. **(G)** Liver beta-hydroxybutyrate concentrations from control fed or overnight fasted mice Data presented as mean ± SD from *n* = 3 independent experiments. Statistical significance was calculated using two-tailed unpaired Student’s *t*-test.

**Supplement Figure 5.**
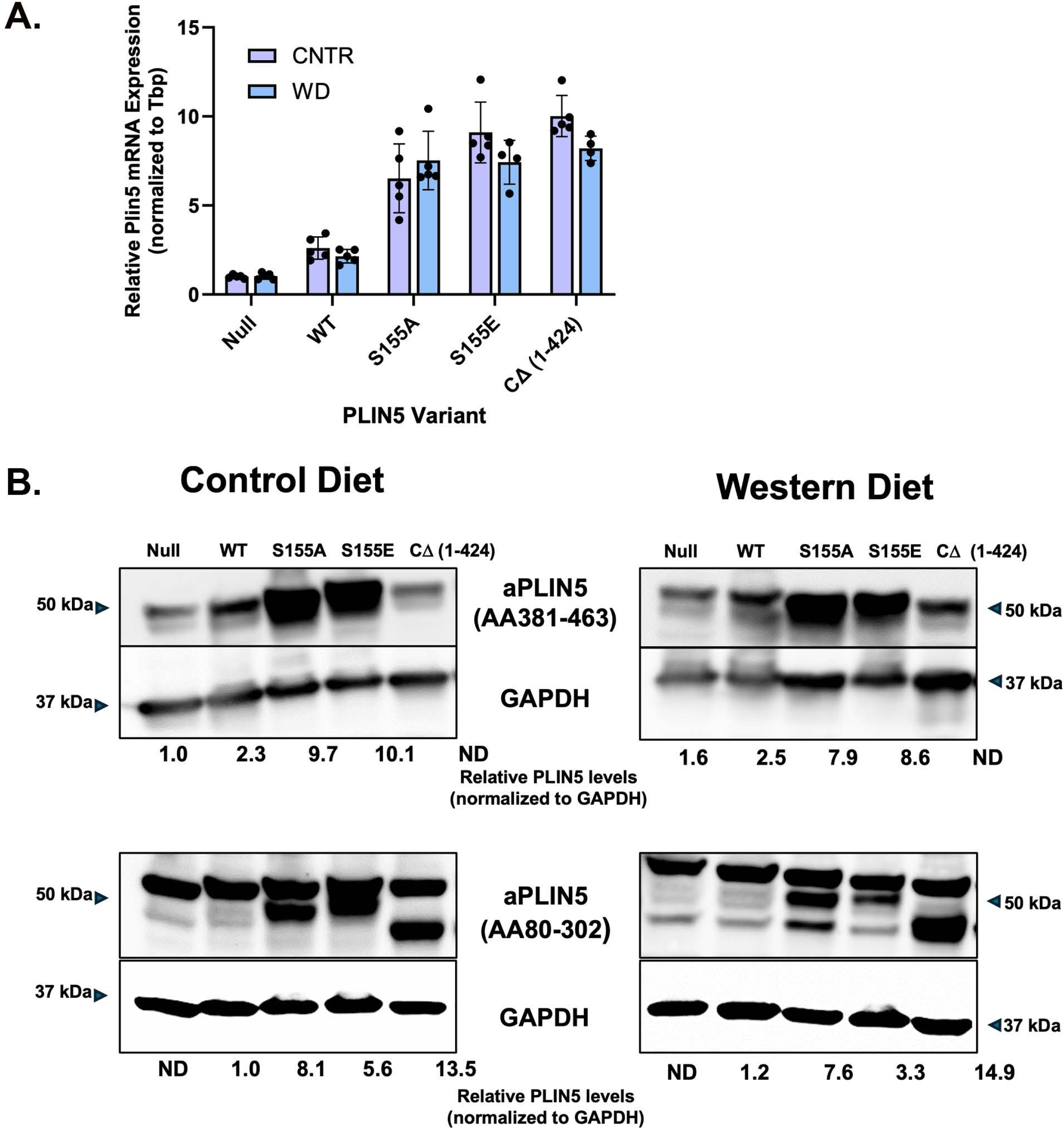
*In vivo* overexpression of PLIN5. **(A)** Relative Plin5 mRNA expression in mouse livers is presented as the mean ± SEM from n=5 except for WD mice expressing S155E and CΔ (1-424) where n=4. **(B)** Representative immunoblot and quantification of PLIN5 protein levels from one mouse per group. An antibody generated against the human PLIN5 carboxy terminus (upper panel) detects endogenous and exogenous full-length PLIN5 but does not detect the CΔ (1-424) variant. PLIN5 levels are relative to endogenous in the Null CNTR mouse. An antibody generated against amino acids within the amino terminus of human PLIN5 (lower panel) detects all forms of exogenous PLIN5 but does not detect endogenous levels in Null CNTR mice. PLIN5 levels are relative to WT exogenous levels in CNTR mice. Full-length PLIN5 migrates at 50 kDa and the CΔ (1-424) PLIN5 migrates slightly below 50 kDa. An intense non-specific >50kDa band is present in all samples and a <50kDa band of undetermined origin is seen in WD-fed mouse samples.

**Supplement Figure 6.**
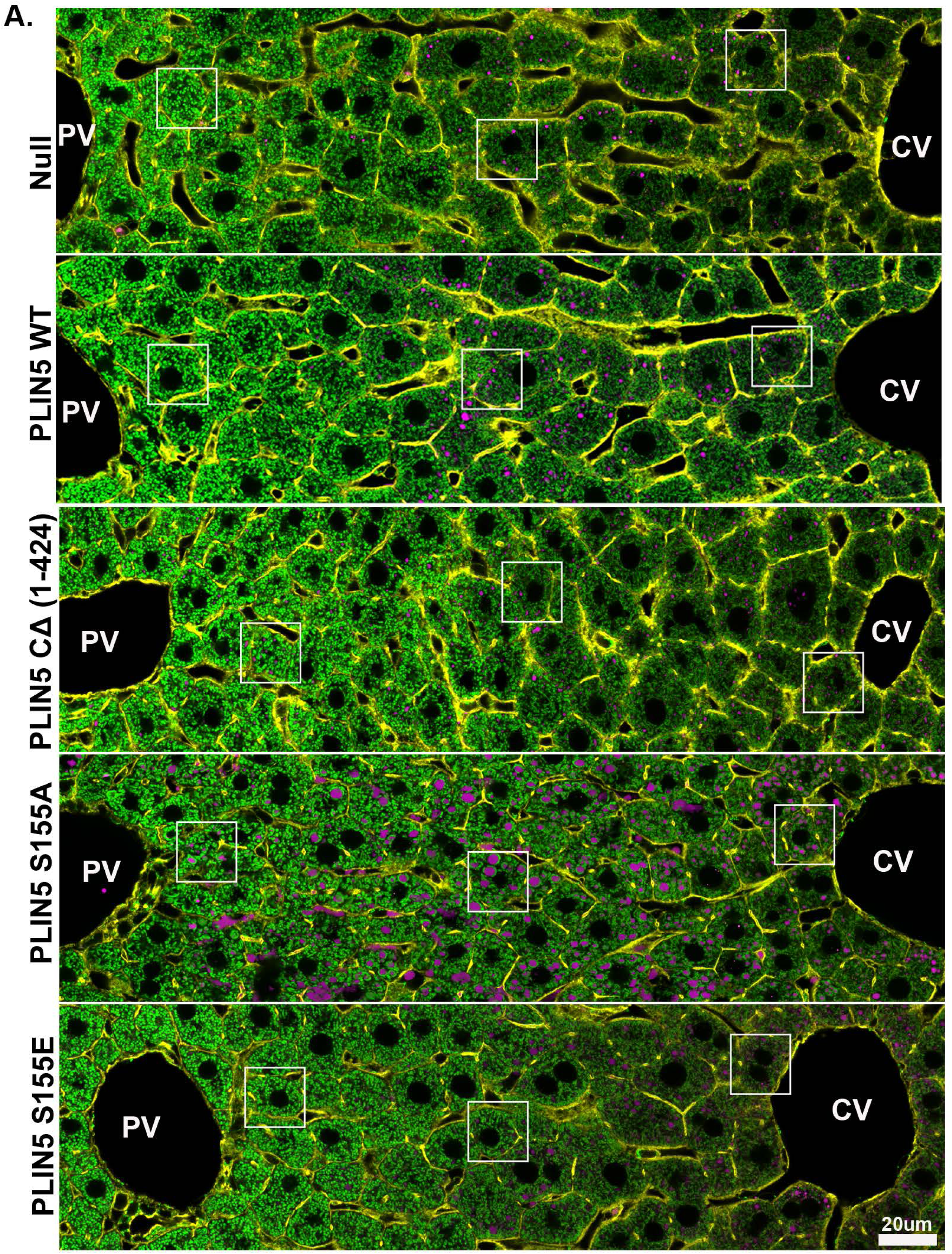
*In vivo* remodeling of mitochondria-LDs interactions. **(A)** Representative confocal images of liver sections from control (CNTR) diet-fed mice overexpressing PLIN5 variants. Images show mitochondria (green), LDs (magenta), and actin (yellow). PV: portal vein; CV: central vein. WT: wild type.

**Supplement Figure 7.**
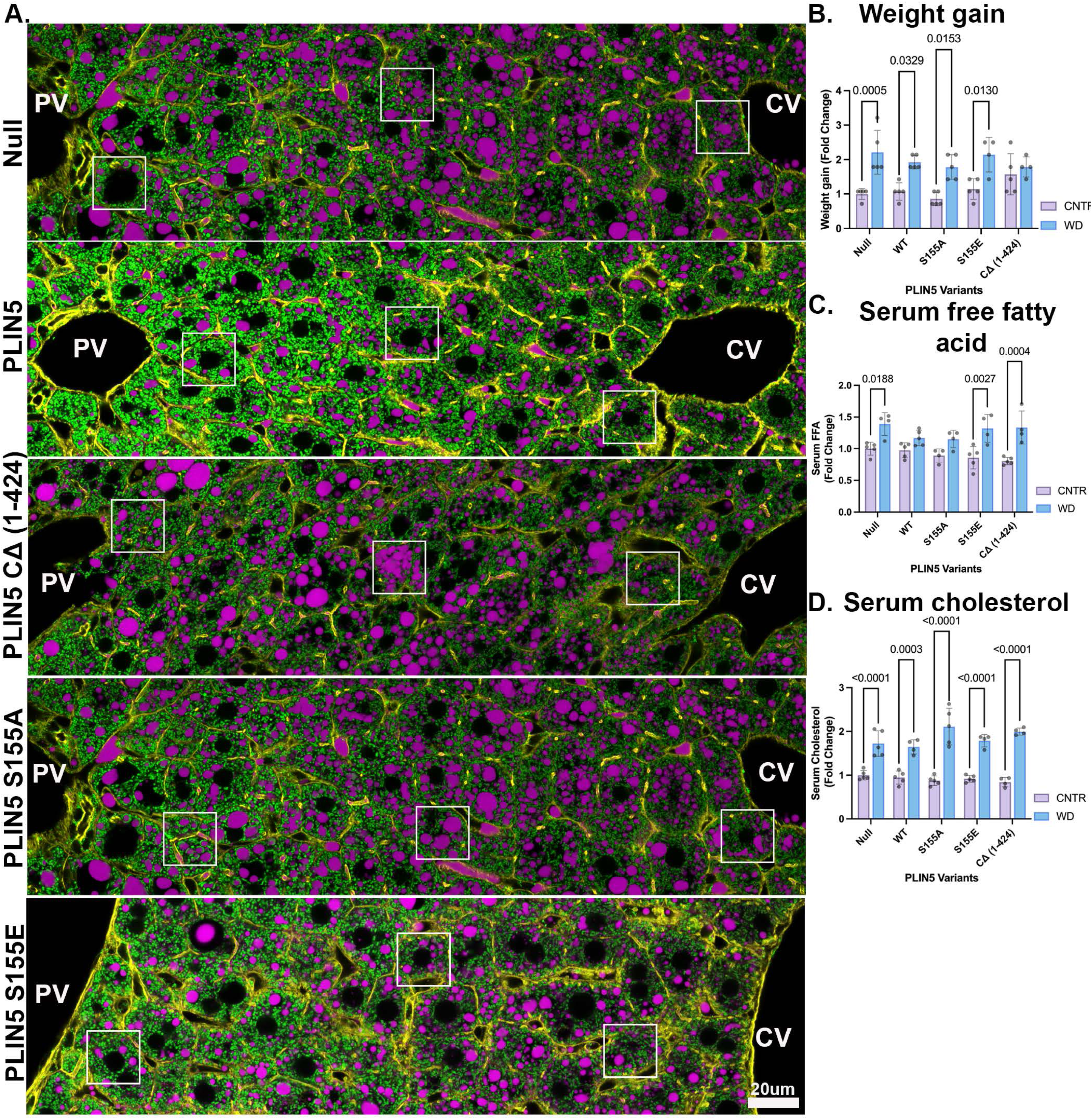
Overexpression of PLIN5 variants in WD-fed mice. **(A)** Representative confocal images of liver sections from Western Diet (WD)-fed mice overexpressing PLIN5 variants. Images show mitochondria (green), LDs (magenta), and actin (yellow). **(B)** Weight gain in mice overexpressing PLIN5 variants and fed CNTR or WD. **(C)** Serum fatty acid levels and **(D)** Serum cholesterol levels in mice overexpressing PLIN5 variants and fed CNTR or WD. Statistical significance was calculated with two-way ANOVA. Data presented as mean ± SD from *n* = 5 independent experiments. PV: portal vein; CV: central vein; WT: wild type. CNTR: control; WD: western diet.

## Materials and methods

### Animal experiments

Experiments were approved by the Institutional Animal Care and Use Committee of the National Cancer Institute and in compliance with the Guide for the Care and Use of Laboratory Animals (National Institutes of Health publication 86-23, revised 1985). All experiments were conducted on ad libitum-fed on normal chow diet (NIH-31 Open Formula) or western diet (TD.120528, Envigo), four to ten weeks old C57BL/6J (strain# 000664) or mtDendra2 excised ^14^ (photo- activatable mitochondria; strain# 018397) male mice obtained from Jackson Laboratories.

### Intra-cardiac fixation, tissue processing, and immunofluorescence

Mice were anesthetized with 250 mg/kg Xylazine and 50 mg/kg Ketamine (diluted in saline) injected intraperitoneally (i.p.). The liver was fixed by transcardial perfusion of ice-cold PBS for 2 min followed by ice-cold 4% paraformaldehyde (PFA) in PBS at a rate of 5 ml/min. Livers were harvested and stored in 4% PFA in PBS overnight and processed in a sucrose gradient before embedding in OCT (Tissue-Tek). Blocks were kept at -80 °C until 10 μm thick slices were made with a cryostat and slides prepared. Slides were stored at -80 °C until thawed, rehydrated, and blocked with 0.1% Triton X-100 and 10% FBS in PBS for 1 h at room temperature. Next, slides were incubated with primary antibody at 4 °C overnight. The following day, the slides were washed three times, 15 min each, and then incubated with a secondary antibody for 1 h at room temperature. After three 15 min washes, slides were mounted with Fluoromount-G and a coverslip (#1).

### Confocal microscopy and scPhenomics

Tile scans and z stacks were acquired using a Leica SP8 inverted confocal laser scanning microscope using a 63x oil objective and a 1.4 numerical aperture.

Whole lobule images were analyzed using a custom Python script (Python v. 3.10) using the deep-learning Cellpose package for segmentation ^15^. Individual hepatocytes were identified by first creating a combination channel of actin, and Gaussian blurred mitochondria. We applied a custom-trained Cellpose model to this combination channel to generate hepatocyte segments. The central and portal veins were manually segmented, and the distance between the central and portal veins was calculated. Segmented hepatocyte distance to the central vein was normalized to the central-portal vein distance and then binned into 12 regions between the portal and central vein.

Mitochondria and lipid droplets were segmented using their own custom-trained Cellpose models. Lipid droplet segments were also subjected to an intensity threshold to remove false positive segments. The mitochondria and lipid droplet segments were used to quantify organelle counts per cell, densities, mitochondrial and lipid fluorescence intensities, as well as geometric parameters, including area, circularity, and eccentricity, using the region props function from scikit-image ^42^. The shortest mitochondria to lipid droplet distance and percent mitochondria/lipid droplet overlap were quantified. For 3D evaluation, surfaces were generated using Imaris (Bitplane 9.9.0) for mitochondria, lipid droplets, and the whole cell. Geometrical measurements were exported into Excel.

### Histology

Liver tissues were also used for independent histological analysis of liver steatosis with hematoxylin and eosin (H&E) by Histoserv (Germantown, MD)

### Liver Perfusion and hepatocyte isolation

Livers of anesthetized mice were perfused by inserting a 22-gauge syringe into the portal vein and delivering 25 ml of pre-warmed (37 °C) perfusion buffer [Krebs-Henseleit buffer (Sigma) with 0.5 μM EDTA], followed by 50 ml collagenase A buffer [Krebs-Henseleit buffer, with 0.1 mM Ca2Cl and 0.4 mg/ml collagenase A (Sigma)]. After perfusion, livers were transferred into a Petri dish, flooded with cold PBS and hepatocytes gently released using forceps. Dissociated cells were collected and filtered through a 100 μm cell strainer. Cells were centrifuged at 50 rpm for 5 min at 4 °C to obtain a hepatocyte enriched pellet. Pellets were resuspended in cold PBS, filtered through a 40 μm cell strainer and centrifuged at 50 rpm for 5 min at 4 °C. Percoll diluted in 10x Hank’s buffer (Sigma) was added to cell suspension and centrifuged at 50 rpm for 5 min at 4 °C. and the supernatant containing dead cells was removed by aspiration. The pellet was resuspended in cold PBS and cell number determined.

### Fluorescence-activated cell sorting (FACS) of PP and PC populations

Cell sorting was performed on a FACSAria Fusion cell sorter (BD Biosciences). Forward and side light scatter was used to distinguish cells from debris and to identify single cells. Zombie-Green fixable live dead dye (1:500, BioLegend) was used to discriminate live and dead cells. The PP and PC populations were identified using anti-E-cadherin-PE and anti-CD73-APC antibody staining (1:150; BioLegend). The PP- and PC-specific gates were set after spectral compensation using appropriate fluorescent minus (FMO) controls.

### Protein Digestion and TMT labeling

Cells were lysed in 50 mM HEPES, pH 8.0, 8M urea and 10% methanol followed by sonication. Lysates were clarified by centrifugation and protein concentration was quantified using BCA protein estimation kit (Thermo Fisher). A 250 ug was alkylated and digested by incubating overnight at 37oC in trypsin at a ratio of 1:50 (Promega). Digestion was acidified by adding formic acid (FA) to a final concentration of 1% and desalted using Pierce peptide desalting columns according to the manufacturer protocol. Peptides were eluted from the column using 50% ACN/0.1% FA, dried in a speedvac and kept frozen in -20oC for further analysis.

For TMT labeling 125 ug of each sample was reconstituted in 50 ?l of 50 mM HEPES, pH 8.0, 500 ug of TMTpro-16plex label (Thermo Fisher) in 100% ACN. After incubating the mixture for 1 h at room temperature with occasional mixing, the reaction was terminated by adding 8 ?l of 5% hydroxylamine. As there were more than 16 samples, we generated a pool sample consisting of equal amounts of lysate from each condition and TMT labeled. The pool was added to each TMT experiment. The TMT-labeled peptides were pooled and dried in a speedvac. The samples were desalted, and excess TMT label removed using peptide desalting columns (Thermo Fisher). A 100 ug aliquot of labeled peptide mixture was fractionated using high pH reversed phase.

### High pH reversed-phase fractionation

The first-dimensional separation of peptides was performed using a Waters Acquity UPLC system coupled with a fluorescence detector (Waters) using a 150 mm x 3.0 mm Xbridge Peptide BEMTM 2.5 μm C18 column (Waters) operating at 0.35 ml/min. The dried peptides were reconstituted in 100 μl of mobile phase A solvent (3 mM ammonium bicarbonate, pH 8.0). Mobile phase B was 100% acetonitrile (Thermo Fisher). The column was washed with mobile phase A for 10 min followed by gradient elution 0- 50% B (10-60 min) and 50-75 %B (60-70 min). Fractions were collected every minute. These 60 fractions were pooled into 24 fractions. Fractions were vacuum centrifuged, and lyophilized fractions were stored at -80°C until analysis by mass spectrometry.

### Mass Spectrometry acquisition

The lyophilized peptide fractions were reconstituted in 0.1% TFA and subjected to nanoflow liquid chromatography (Thermo UltimateTM 3000RSLC nano LC system, Thermo Fisher) coupled to an Orbitrap Eclipse mass spectrometer (Thermo Fisher). Peptides were separated using a low pH gradient using a 5-50% ACN over 120 min in a mobile phase containing 0.1% formic acid at 300 nl/min flow rate. MS scans were performed in the Orbitrap analyzer at a resolution of 120,000 with an ion accumulation target set at 4e5 and max IT set at 50ms over a mass range of 400-1600 m/z. Ions with a determined charge state between 2 and 5 were selected for MS2 scans in the ion trap with CID fragmentation (Turbo; NCE 35%; maximum injection time 35 ms; AGC 1 × 104). The spectra were searched using the Real Time Search Node in the tune file using mouse Uniprot database using Comet search algorithm with TMT16 plex (304.2071Da) set as a static modification of lysine and the N-termini of the peptide. Carbamidomethylation of cysteine residues (+57.0214 Da) was set as a static modification, while oxidation of methionine residues (+15.9949 Da) was set up as a dynamic modification. For the selected peptide, an SPS–MS3 scan was performed using up to 10 b- and y-type fragment ions as precursors in an Orbitrap at 50,000 resolution with a normalized AGC set at 500 followed by maximum injection time set as “Auto” with a normalized collision energy setting of 65.

## Data Analysis

Acquired MS/MS spectra were searched against the mouse Uniprot protein database along with a contaminant protein database using SEQUEST in the Proteome Discoverer 2.4 software (Thermo Fisher). The precursor ion tolerance was set at 10 ppm and the fragment ions tolerance was set at 0.02 Da, with methionine oxidation included as a dynamic modification. Carbamidomethylation of cysteine residues and TMTpro16 plex (304.2071 Da) were set as static modification of lysine and the N-terminus of the peptide. Trypsin was specified as the proteolytic enzyme, with up to 2 missed cleavage sites allowed. Searches used a reverse sequence decoy strategy to control for the false peptide discovery and identifications were validated using the Percolator algorithm in Proteome Discoverer 2.4.

Reporter ion intensities were adjusted to correct for lot-specific impurities according to the manufacturer specification and the abundances of the proteins were quantified using the summation of the reporter ions for all identified peptides. A two-step normalization procedure was applied on a 24-plex TMT experiment [ 2 TMT experiments with 12 samples each]. First, the reporter abundances were normalized across all the channels to account for equal peptide loading. Secondly, the intensity from the pooled sample was used to normalize the batch effect from the multiple TMT experiments. Samples were further normalized using the voom algorithms and quantile normalization from the Limma R package (v3.40.6). Differential peptide expression analysis was performed using Limma, where zonation was determined by the p-value (0.05) and/or FC (?1.2); and pathway enrichment analysis was performed using Fisher’s Exact test with GO database. Graphpad Prism 9 and Biorender was used for visualization.

### Seahorse assay

A Seahorse XFe96 Analyzer and XF Mito Stress Test Kit (Agilent Technologies) was used to measure the oxygen consumption rate (OCR) and extracellular acidification rate (ECAR) of PP and PC sorted cells. A day prior to the assay, the cartridge sensor was hydrated overnight with Seahorse Bioscience Calibration Solution at 37 °C without CO2. After sorting, cells were seeded in a 96-well seahorse plate at a density of 5.0 × 104 cells/well and allowed to adhere for 2 h. After 2 h, cell media was replaced with serum-free Seahorse XF Base media containing glucose, pyruvate, and glutamine (pH 7.4), and the cells were incubated at 37 °C in a non-CO2 incubator for 1 h. OCR and ECAR were measured after the injection of oligomycin (1.5 μM), FCCP (1 μM), and rotenone/antimycin (0.5 μM). In addition to XF Mito Stress Test Kit, XF Mito Fuel kit was used together includes three inhibitors, UK5099 (2 μM), BPTES (3 μM) and etomoxir (4 μM) to measure the dependency of cells on the three major mitochondrial fuels; glucose, glutamine and long- chain fatty acids, respectively. Four independent experiments were conducted with six to eight replicates per population and treatment condition per experiment. All data was normalized to BCA (Thermo Fisher). Measurements for basal respiration and maximum respiration capacity were presented as a percentage normalized to PP fed.

### Overexpression of PLIN5 variants in mice

Four-week-old C57BL/6 J male mice and mtDendra2 excised male mice were housed five per cage in a facility (12 h light/dark cycle) and fed either a control chow diet (NIH-31 Open Formula) or Western Diet (WD; TD.120528, Envigo) for two weeks. Mice were then injected via tail vein with an AAV8-ABLp vector: Null, mPlin5 WT, S155A, S155E, CΔ (1-424), at a titer of 4.5× 1011 GC (Vector Biolabs, Philadelphia, PA). After AAV8 transduction, mice were fed the control chow diet or WD for four weeks. At the end of the experiment, mice serum and liver tissue were collected.

### Serum analysis

Kits were used according to manufacturer’s instructions to measure β-hydroxybutyrate (Abcam), free fatty acid FUJIFILM (Wako Diagnostics), triglycerides (Pointe Scientific), cholesterol (Thermo Scientific), insulin (Crystal Chem Inc).

### Blood Glucose

Blood was collected from the tail vein, and glucose was measured using testing strips and monitor (Ascencia Diabetes Care).

### Triglyceride (TG), Malondialdehyde (MDA), and Glutathione measurements

Pulverized liver tissue samples were lysed, and the supernatant was collected after centrifugation at 13,000 rpm for 10 minutes. Triglyceride levels were measured with a Colorimetric Assay Kit (Elabscience), Cellular Malondialdehyde (MDA) levels were measured with a Lipid Peroxidation (MDA) Assay Kit (Abcam), and total cellular glutathione (GSH) levels were estimated through the GSH Glo™ Assay Kit (Promega). Each kit was used per the manufacturer’s instructions, and results normalized to protein amount, and absolute numbers converted to fold change.

### Western blot analysis

Proteins from primary hepatocytes or pulverized liver were extracted by homogenization in RIPA lysis buffer (150 mM NaCl, 0.1% Triton X-100, 0.5% sodium deoxycholate, 0.1% sodium dodecyl sulfate, and 50 mM Tris HCl pH 8.0) containing EDTA, PMSF, and Halt Inhibitor Cocktail (Thermo Fisher), followed by centrifugation at 13,000 rpm at 4 °C for 30 min. Protein concentration was determined using a Pierce™ BCA Protein Assay Kit (Thermo Fisher). Lysates were boiled at 95 °C for 5 min, and 5-10 μg aliquots were fractionated by SDS-PAGE and then transferred to nitrocellulose 0.45 μm membrane (Bio-Rad). Membranes were blocked for 1 h at room temperature in 5% BSA in 1x Tris-buffered saline + 0.1% Tween 20 (TBST) then incubated with primary antibody diluted in 5% BSA in 1x TBST at 4 °C overnight. Membranes were washed three times with TBST then incubated in the secondary antibody 1 h at room temperature. Membranes were washed three times with TBST, then Clarity ECL western blot substrate solution (Bio-Rad) was applied for detection and imaged using the ChemiDoc Imaging System (Bio-Rad). Bands were quantified with ImageJ.

### RNA isolation and Quantitative PCR

RNA was isolated from pulverized liver tissue using the TissueLyser LT and RNeasy mini kit (Qiagen). A High-Capacity RNA to cDNA kit (Thermo Fisher) was used to synthesize random-primed cDNA from 2 μg DNAse treated RNA. Real time PCR was conducted in 384 well-plates using a ViiA7 Real-Time PCR system (Applied Biosystems). Singleplex reactions (5 μl) containing a FAM-MGB expression assay for Plin5 (Mm00508854_m1) or Tbp control (Mm01277041_m1) (Thermo Fisher) were performed using cDNA synthesized from 8 ng RNA and 1× Fast Advanced Master Mix (Thermo-without Amp Erase UNG) were performed in triplicate. The comparative Ct method (delta, delta Ct) was used to determine relative expression normalized to Tbp (Applied Biosystems® ViiA™ 7Real-Time PCR System Getting Started Guides).

**Supplement table 1:**
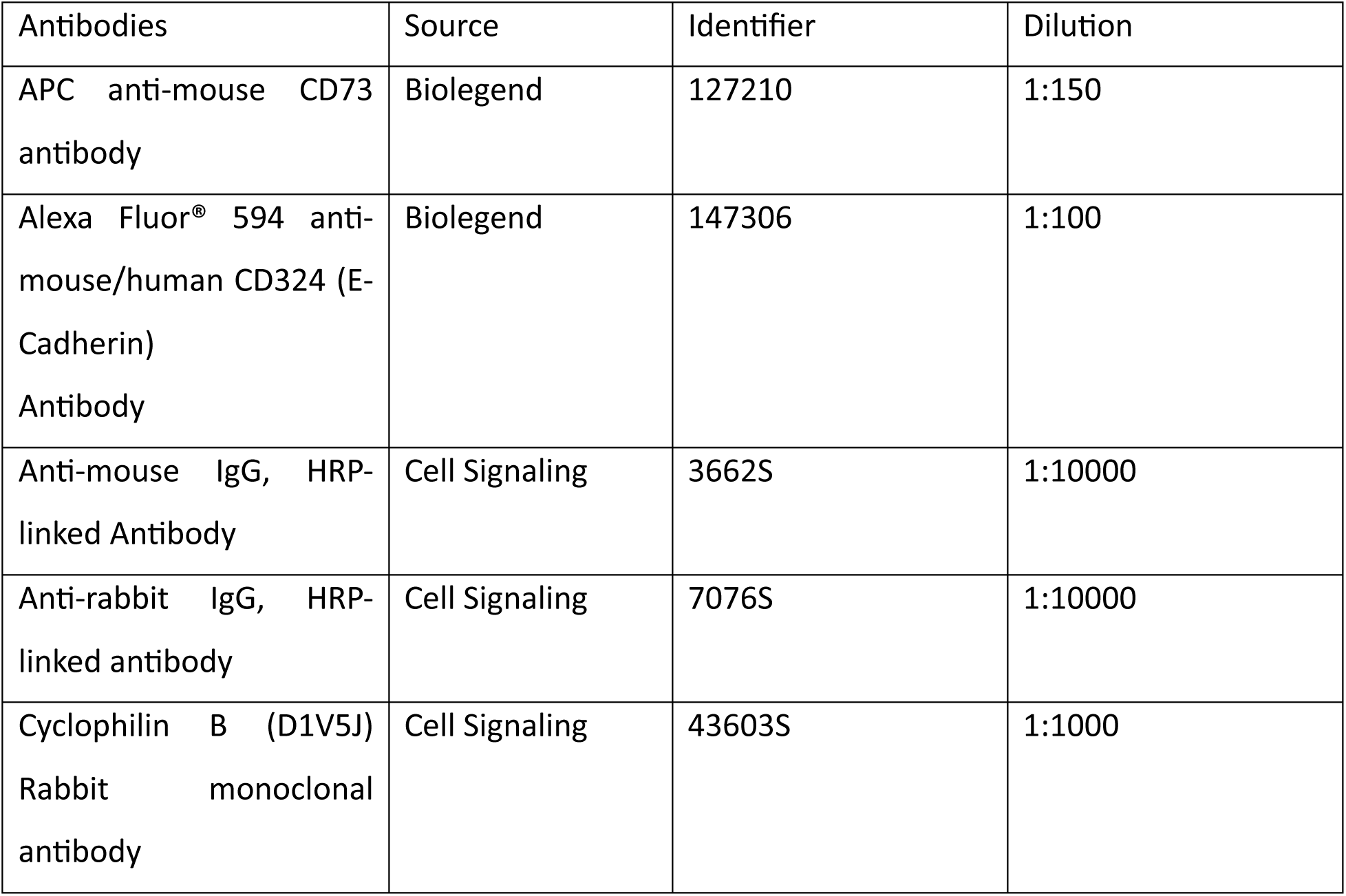
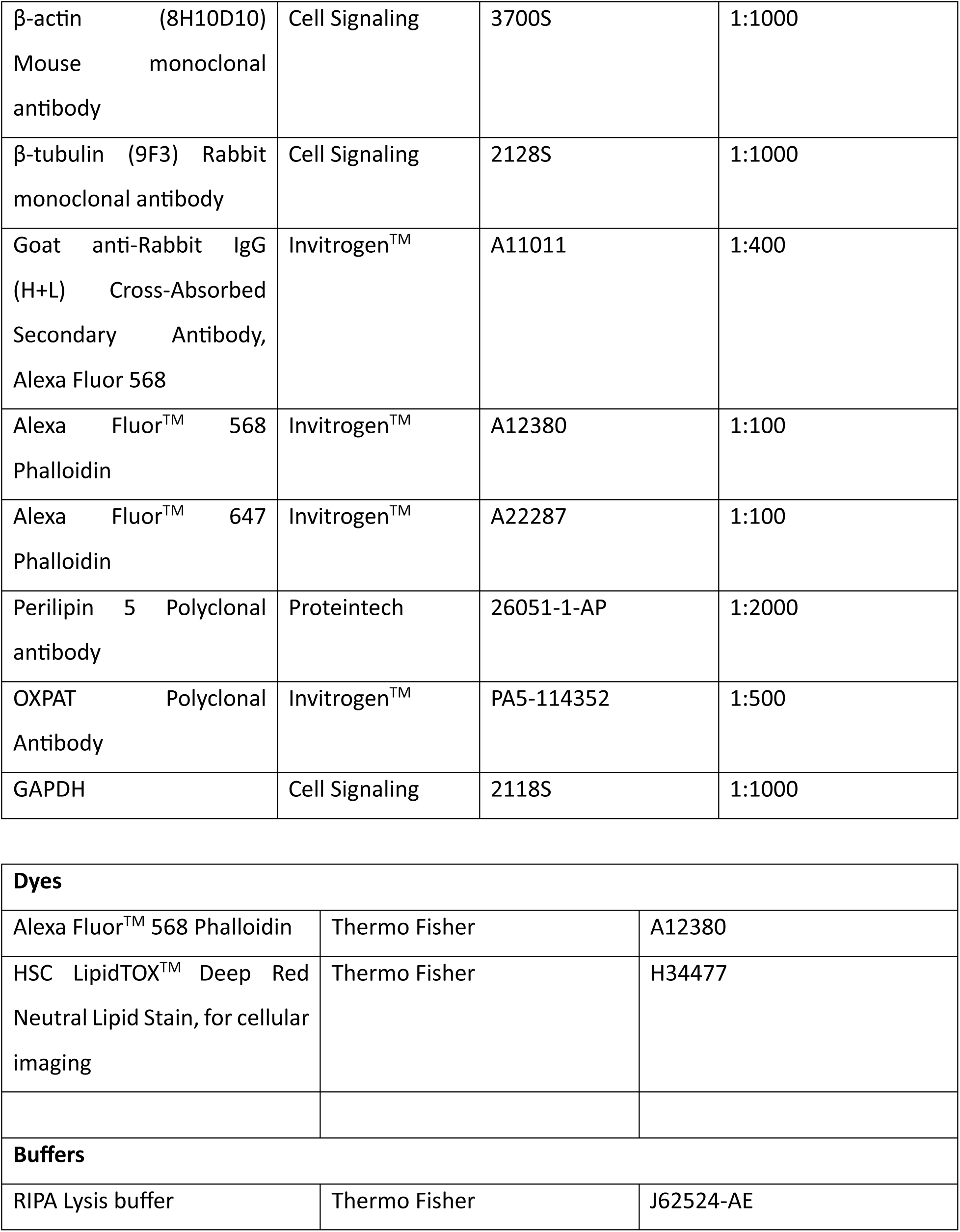
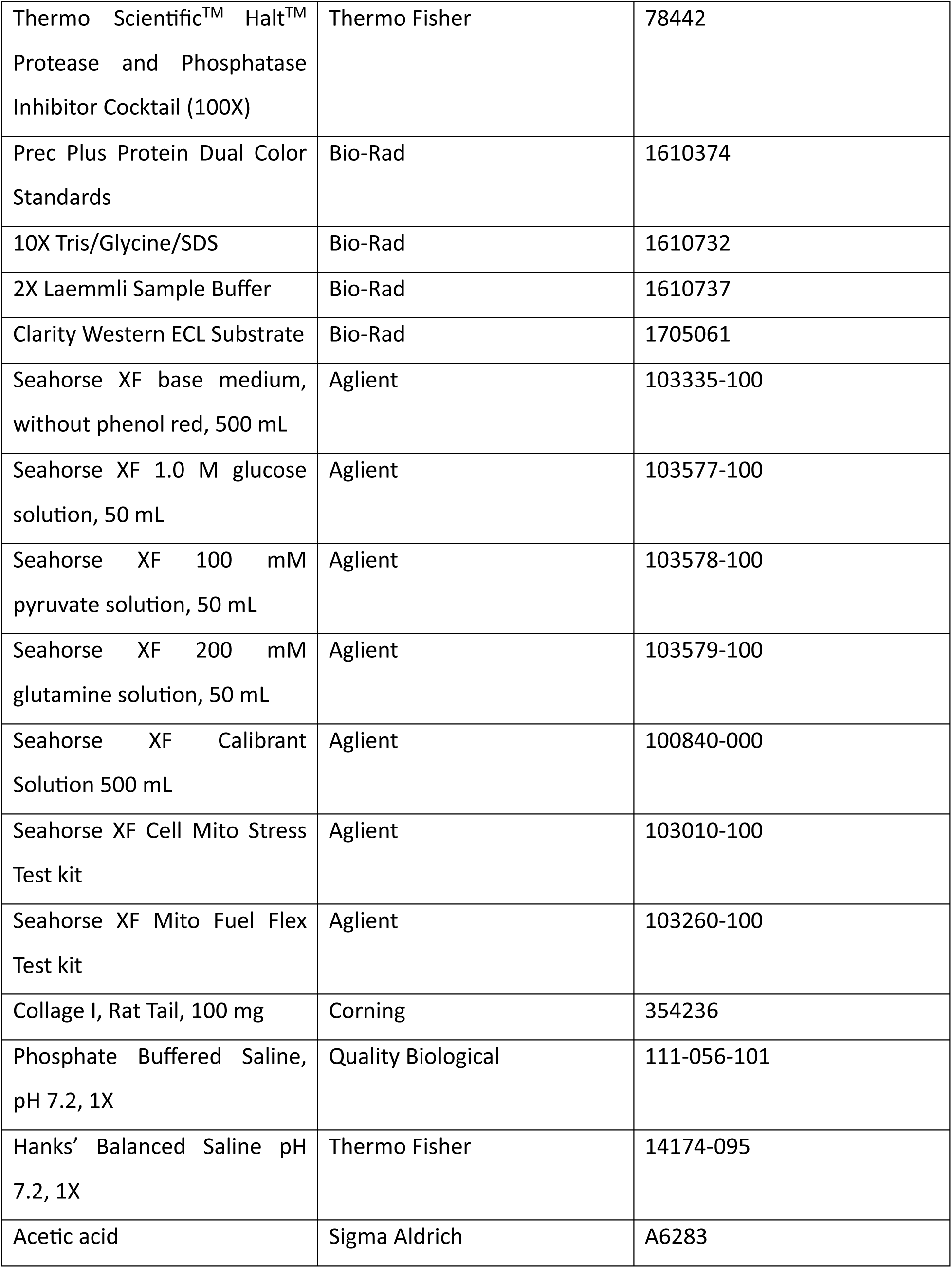
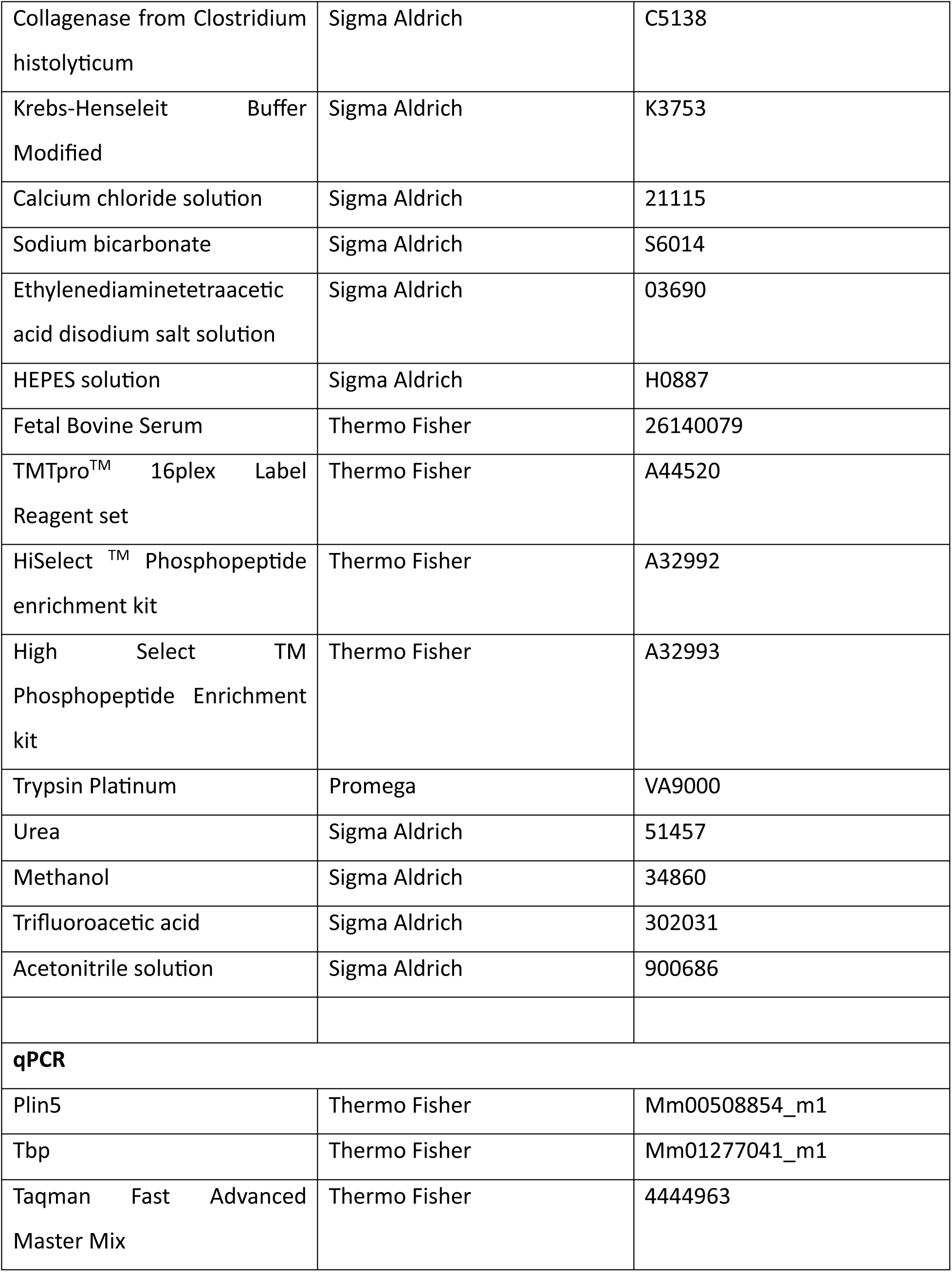
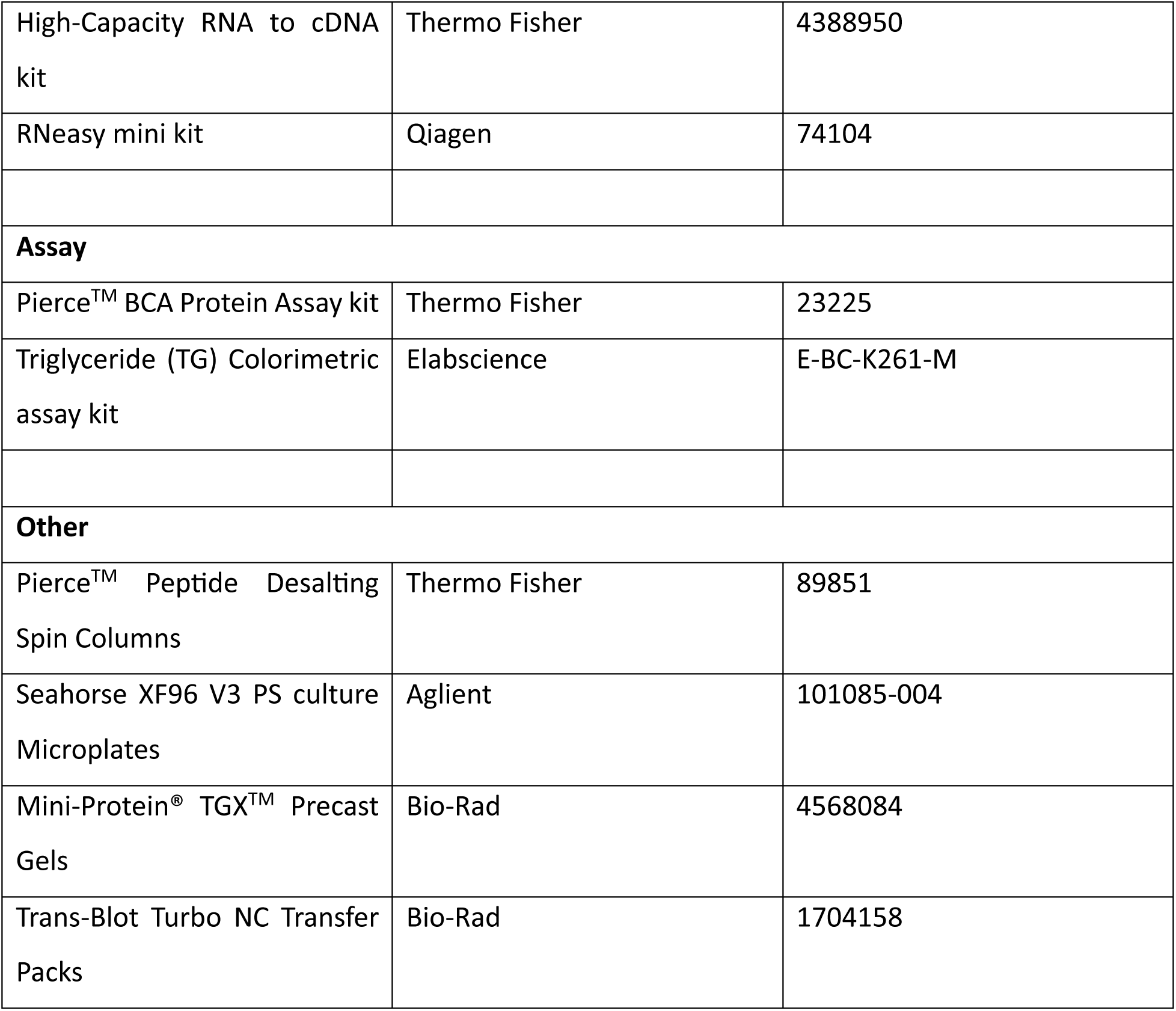

| Antibodies | Source | Identifier | Dilution |
| --- | --- | --- | --- |
| APC anti-mouse CD73 antibody | Biolegend | 127210 | 1:150 |
| Alexa Fluor® 594 anti-mouse/human CD324 (E-Cadherin) Antibody | Biolegend | 147306 | 1:100 |
| Anti-mouse IgG, HRP-linked Antibody | Cell Signaling | 3662S | 1:10000 |
| Anti-rabbit IgG, HRP-linked antibody | Cell Signaling | 7076S | 1:10000 |
| Cyclophilin B (D1V5J) Rabbit monoclonal antibody | Cell Signaling | 43603S | 1:1000 |
| β-actin (8H10D10)<br>Mouse monoclonal<br>antibody | Cell Signaling | 3700S | 1:1000 |
| β-tubulin (9F3) Rabbit<br>monoclonal antibody | Cell Signaling | 2128S | 1:1000 |
| Goat anti-Rabbit IgG<br>(H+L) Cross-Absorbed<br>Secondary Antibody,<br>Alexa Fluor 568 | Invitrogen™ | A11011 | 1:400 |
| Alexa Fluor™ 568<br>Phalloidin | Invitrogen™ | A12380 | 1:100 |
| Alexa Fluor™ 647<br>Phalloidin | Invitrogen™ | A22287 | 1:100 |
| Perilipin 5 Polyclonal<br>antibody | Proteintech | 26051-1-AP | 1:2000 |
| OXPAT Polyclonal<br>Antibody | Invitrogen™ | PA5-114352 | 1:500 |
| GAPDH | Cell Signaling | 2118S | 1:1000 |

|  |  |  |
| --- | --- | --- |
| **Dyes** |  |  |
| Alexa Fluor™ 568 Phalloidin | Thermo Fisher | A12380 |
| HSC LipidTOX™ Deep Red<br>Neutral Lipid Stain, for cellular<br>imaging | Thermo Fisher | H34477 |
| **Buffers** |  |  |
| RIPA Lysis buffer | Thermo Fisher | J62524-AE |
| Thermo Scientific™ Halt™ Protease and Phosphatase Inhibitor Cocktail (100X) | Thermo Fisher | 78442 |
| Prec Plus Protein Dual Color Standards | Bio-Rad | 1610374 |
| 10X Tris/Glycine/SDS | Bio-Rad | 1610732 |
| 2X Laemmli Sample Buffer | Bio-Rad | 1610737 |
| Clarity Western ECL Substrate | Bio-Rad | 1705061 |
| Seahorse XF base medium, without phenol red, 500 mL | Aglient | 103335-100 |
| Seahorse XF 1.0 M glucose solution, 50 mL | Aglient | 103577-100 |
| Seahorse XF 100 mM pyruvate solution, 50 mL | Aglient | 103578-100 |
| Seahorse XF 200 mM glutamine solution, 50 mL | Aglient | 103579-100 |
| Seahorse XF Calibrant Solution 500 mL | Aglient | 100840-000 |
| Seahorse XF Cell Mito Stress Test kit | Aglient | 103010-100 |
| Seahorse XF Mito Fuel Flex Test kit | Aglient | 103260-100 |
| Collage I, Rat Tail, 100 mg | Corning | 354236 |
| Phosphate Buffered Saline, pH 7.2, 1X | Quality Biological | 111-056-101 |
| Hanks' Balanced Saline pH 7.2, 1X | Thermo Fisher | 14174-095 |
| Acetic acid | Sigma Aldrich | A6283 |
| Collagenase from Clostridium histolyticum | Sigma Aldrich | C5138 |
| Krebs-Henseleit Buffer Modified | Sigma Aldrich | K3753 |
| Calcium chloride solution | Sigma Aldrich | 21115 |
| Sodium bicarbonate | Sigma Aldrich | S6014 |
| Ethylenediaminetetraacetic acid disodium salt solution | Sigma Aldrich | 03690 |
| HEPES solution | Sigma Aldrich | H0887 |
| Fetal Bovine Serum | Thermo Fisher | 26140079 |
| TMTpro™ 16plex Label Reagent set | Thermo Fisher | A44520 |
| HiSelect™ Phosphopeptide enrichment kit | Thermo Fisher | A32992 |
| High Select™ Phosphopeptide Enrichment kit | Thermo Fisher | A32993 |
| Trypsin Platinum | Promega | VA9000 |
| Urea | Sigma Aldrich | 51457 |
| Methanol | Sigma Aldrich | 34860 |
| Trifluoroacetic acid | Sigma Aldrich | 302031 |
| Acetonitrile solution | Sigma Aldrich | 900686 |
| **qPCR** |  |  |
| Plin5 | Thermo Fisher | Mm00508854_m1 |
| Tbp | Thermo Fisher | Mm01277041_m1 |
| Taqman Fast Advanced Master Mix | Thermo Fisher | 4444963 |
| High-Capacity RNA to cDNA kit | Thermo Fisher | 4388950 |
| RNeasy mini kit | Qiagen | 74104 |
| **Assay** |  |  |
| Pierce™ BCA Protein Assay kit | Thermo Fisher | 23225 |
| Triglyceride (TG) Colorimetric assay kit | Elabscience | E-BC-K261-M |
| **Other** |  |  |
| Pierce™ Peptide Desalting Spin Columns | Thermo Fisher | 89851 |
| Seahorse XF96 V3 PS culture Microplates | Aglient | 101085-004 |
| Mini-Protein® TGX™ Precast Gels | Bio-Rad | 4568084 |
| Trans-Blot Turbo NC Transfer Packs | Bio-Rad | 1704158 |

